# Fabricating microfluidic co-cultures of immortalised cell lines uncovers robust design principles for the simultaneous formation of patterned, vascularised, and stem cell-derived adipose tissue

**DOI:** 10.1101/2025.01.22.634386

**Authors:** Ashley R. Murphy, Rose Ann Franco, Mark C. Allenby

## Abstract

*In vitro* culture processes supporting the simultaneous formation of vessel networks alongside differentiation towards mature parenchymal tissue have numerous clinical and agricultural applications but remain unrealised due to contrasting culture requirements. Of specific interest is lab-grown vascularised adipose tissue to study metabolic syndrome, diabetes, obesity, cardiovascular diseases, and to advance cultivated meat technologies. We report a microfluidic 3D hydrogel culture device capable of supporting live-imaging of fluorescent reporter cell lines and generating counter-current gradients of vasculogenic and adipogenic growth factors. for the first time, we report experimental conditions capable of reproducibly forming diverse microvascular networks from telomerase immortalised endothelial and mesenchymal stem cells in both 2D and 3D hydrogel-embedded cultures. Using our novel microfluidic culture design, we demonstrate the generation of growth factor environments which support the 3D co-formation of integrated robust microvascular networks and lipid-producing adipocytes after 31 days gradient culture. We demonstrate microvascular networks substantially support parenchymal stromal cell differentiation to mature adipose tissue (67.4% lipid coverage), unachieved in avascular cultures (1.86% lipid coverage). We attempt to validate our co-culture model by applying inhibitors of vessel-mediated lipogenesis (spermidine and VO-OHpic), which are demonstrated to be ineffective in our novel human preclinical model.

**Highlights:**

- An optimised co-culture protocol for vasculogenesis of immortalised cell lines.
- Immortalised co-cultures form reproducible microvessel networks up to 31 days.
- Gradient co-culture enables simultaneous MSC adipogenesis and EC vasculogenesis.
- MSC differentiation to dense, lipid-laden adipose tissue relies on EC co-culture.
- Human vascularised adipose tissue formation is unaffected by a murine regulator.

**Graphical Abstract:** 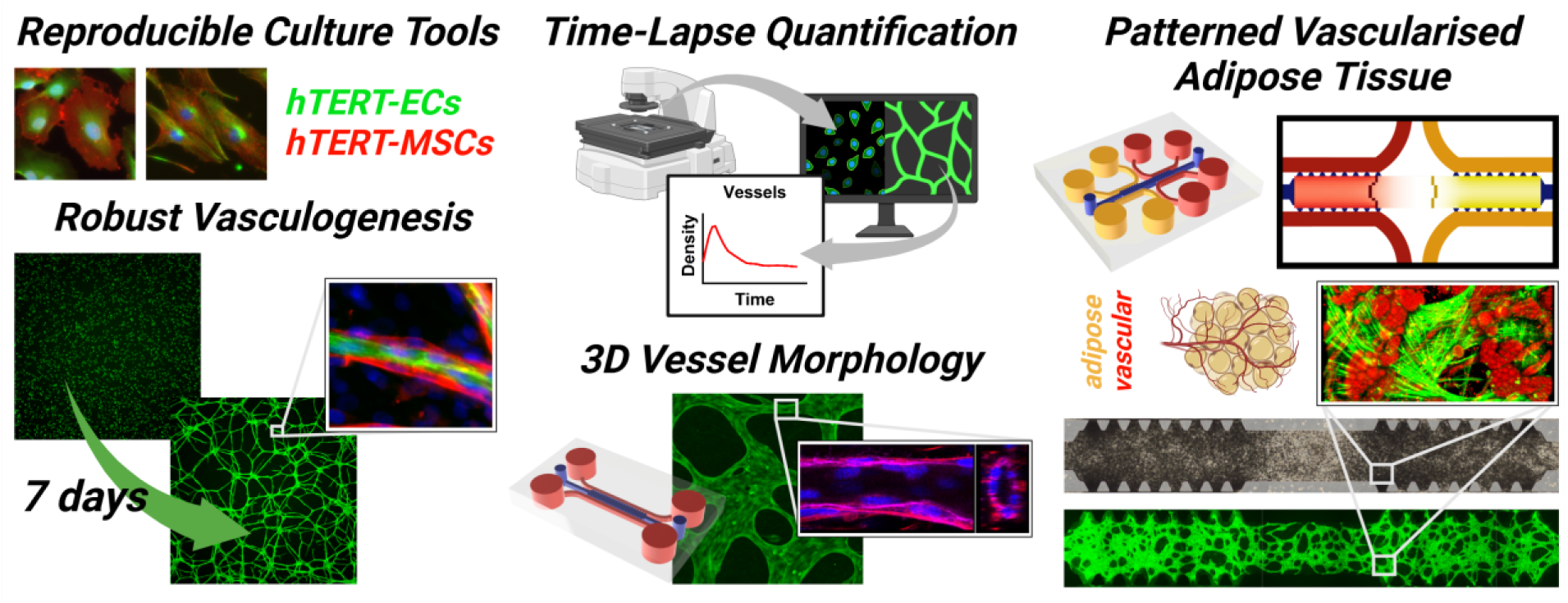

## 1. Introduction

Rapid development in miniaturised tissue and organ-like technology represents remarkable progress towards mimicking the form and function of native biological structures and realising the ultimate goal of restoring damaged tissue or whole organs [1]. However, the limited ability of nutrients and oxygen to diffuse into dense cellular structures can often lead to cell necrosis, inhomogeneous tissue development and limited size [2]. Vasculature is the human bodies inherent plumbing network which supports the transport of nutrients, salts, oxygen, cells and waste. Additionally, endothelium-derived paracrine (angiocrine) signalling, such as anti-inflammatory and pro-inflammatory factors during homeostasis and disease, is critical to parenchymal tissue development, maintenance and repair [3]. More recently, endothelium-secreted small molecules, metabolites and proteins have been demonstrated to regulate critical metabolic processes in parenchymal-specific tissue such as skeletal muscle, the liver and adipose tissue [4–7]. Without functional and physiologically-relevant vascular networks, or the means to actively incorporate host vasculature post-implantation, the development of *in vitro* tissue and organ products will never progress towards clinically relevant scales and physiological complexities [8].

Widespread 2D *in vitro* tissue culture techniques remain limited in supporting the study of pathophysiological contributions by vascular networks on parenchymal tissue [9]. Endothelial cells are typically grown in mono-culture, requiring unique nutrients and growth factors often incompatible with the expansion or differentiation of other cell types [10]. Additionally, 2D cultures deliver over saturated nutrients, growth factors and remove waste products as a batch exchange, leading to homogenous nutrient distributions with limited ability to support endothelial-parenchymal tissue development [10]. In contrast, our body’s complex vascular interactions and heterogenous distributions of nutrients, growth factors, and cells evolve in a highly orchestrated manner and facilitate tissue morphogenesis during development and wound healing [11]. The inability of current *in vitro* models to reproducibly create vascularised parenchymal tissue using basic culture protocols remains a hurdle for the translation of biomanufactured tissue.

Simple partitioned culture systems such as membrane culture inserts are well-used to co-culture two cell types. However, active cell-cell interactions and the controlled establishment of 3D gradients is often not possible. Microfluidic culture devices are increasingly used to establish, control and maintain nutrients gradients for cell and tissue growth [12]. Mesenchymal stem cells (MSCs) have been shown to facilitate endothelial cell vasculogenesis in numerous co-culture models and are additionally multipotent and therefore able to differentiate towards connective tissue lineages such as bone, cartilage and adipose [13]. Utilising the differentiation of resident MSCs in vasculogenic co-cultures has the potential to produce vascularised and mature connective tissues, a goal that remains unrealised due to conflicting media requirements.

White adipose tissue (WAT) is the most abundant adipose tissue in the human body, primarily comprised of 80–90% (v/v) adipocytes with the remainder attributed to vasculature containing endothelial cells, pericytes, fibroblasts, preadipocytes, macrophages and other blood cells [14, 15]. Traditionally, the study of adipose tissue is conducted in monolayer culture, in the absence of these supporting cell types [16], despite WAT vascular networks having critical roles in controlling adipocyte nutrient uptake, facilitating adipogenesis and regulating adipose tissue expansion [17]. Recent 3D co-cultures of adipose progenitors and microvascular cells have been developed; however, these require multi-step differentiation process unable to support the formation microvascular networks alongside differentiating adipose tissue [18, 19].

Simultaneous formation is critical to model adipo-vascular cross-talk, whose dysfunction results in conditions such as metabolic syndrome, diabetes, obesity and cardiovascular diseases [20–25], which highlights the need for physiologically relevant human *in vitro* models of vascularised adipose tissue.

Using bespoke microfluidic device configurations, we establish spatiotemporal counter-current gradients of vasculogenic and adipogenic culture environments across a 3D hydrogel-supported co-culture system. By co-culturing highly reproducible telomerase reverse transcriptase (hTERT) immortalised human adipose-derived mesenchymal stem cells and hTERT immortalised human aortic endothelial cells, we report conditions required to form robust and optimal microvascular networks in 2D and suspended in 3D culture environments. Additionally, we utilise weeklong time-lapse microscopy to extensively quantify and characterise vascular morphogenesis and provide informed observations regarding its mechanism. We explore the ability of gradient culture systems to support the co-formation of microvascular networks and adipogenic differentiation of mesenchymal stem cells and report the enhancing effects of vascular-mediated adipogenesis. We then apply our human preclinical vascularised adipose tissue model to identify the impact of known vasculo-adipo regulators.

## 2. Methods

### 2.1. Microfluidic device design, fabrication and preparation

Microfluidic devices were designed using AutoCAD (Autodesk, Inc.) 3D design software. Silicon master wafters (7-inch diameter) featuring said designs were then prepared by the Australian National Fabrication Facility Queensland Node. Briefly, a UV mask aligner was used to transfer designs onto a photoresist-coated silicon master wafer to a feature height of 200 µm.

Silicon master wafers were treated overnight with the vapour of hexamethyldisilizane and then secured in Petri dishes using Scotch tape. Using SYLGARD 184 Silicone Elastomer Kit (Dow), 10-parts monomer was added with 1-part co-monomer and simultaneously mixed and degassed using a Kurabo KK-V300SS planetary centrifugal mixer and poured onto the master wafer to a height of approximately 12 mm above the wafer. The mixture was then cured at 80°C for a minimum of 1 hour. The cured elastomer was then carefully removed using a scalpel and media reservoirs and cell injection ports were created using 1.25-and 4-mm diameter biopsy punches, respectively. Both elastomer and microscope slides were cleaned with Scotch tape, placed face-up on the stage of a Tergeo Tabletop Plasma Cleaner and treated with oxygen plasma at 35 W for 40 seconds. Elastomer and slide were bonded with direct contact before full adhesion was achieved after approximately 5 minutes.

Microfluidic devices were wet autoclave sterilised at 121°C before being washed of any residue via complete immersion in 70% (w/v) ethanol for a minimum of 1 hour. Ethanol was then completely removed from the devices via vacuum aspiration before the devices were washed in triplicate with sterile distilled water (15230162). Devices were finally dried overnight in a humidified incubator at 37°C in 5% (v/v) CO_2_ in air to remove any remaining water from channels.

### 2.2. Cell culture and imaging

This project was supported by UQ’s Human Research Ethics Committee through approval 2021/HE002698: “tissue culture models for in vitro optimisation of engineered biomaterial scaffolds and bioprocess techniques”, as well as UQ’s Institutional Biosafety Committee through approval IBC/602E/ChemEng/2023: “risk group 2 immortalised cell line activities.”

#### 2.2.1. Routine cell line maintenance

Human telomerase reverse transcriptase (hTERT) immortalized adipose-derived mesenchymal stem cells (hAD-MSCs; ASC52telo, SCRC-4000, American Type Culture Collection ATCC) were routinely maintained as per manufacturer’s instructions in MSC Basal Medium (#500-040) supplemented with Mesenchymal Stem Cell Growth Kit for Adipose and Umbilical-derived MSCs - Low Serum (#500-030), herein referred to as ‘complete MSC media’, and subcultured at a confluence of approximately 80%.

hTERT immortalized green fluorescent protein-expressing human aortic endothelial cells (GFP-hAECs; TeloHAEC-GFP, CRL-4054, ATCC) were routinely maintained as per manufacturers instruction in Vascular Cell Basal Medium (#100-0303) supplemented with Endothelial Cell Growth Kit - VEGF (#100-041) and 33 µM phenol red (#999-001), herein referred to as ‘complete vascular cell media’, and subcultured at a confluence of approximately 80%. Maintenance cultures of both cell types were routinely tested for mycoplasma contamination using MycoStrip® - Mycoplasma Detection Kit (rep-mys-50).

#### 2.2.2. 2D vasculogenesis culture

To induce spontaneous formation of vascular networks from endothelial cells (vasculogenesis), hAD-MSCs were co-cultured with GFP-hAECs in a humidified incubator at 37°C in 5% (v/v) CO_2_ in air. In brief, hAD-MSCs and GFP-hAEC maintenance cultures were washed with room temperature Dulbecco’s phosphate buffered saline solution (DPBS, 14190144, 3 mL/25 cm^2^) before being exposed to 0.05% trypsin-EDTA solution (25300054, 1 mL/25 cm^2^) for approximately 5 minutes, at room temperature for hAD-MSCs, and 37°C for hAECs. Cultures were then neutralised with an equal volume of respective maintenance media, before being centrifuged at 200 × g for 5 minutes at room temperature. Cell pellets were resuspended and counted via trypan blue exclusion method with a Countess 3 Automated Cell Counter (ThermoFisher Scientific). hAD-MSCs and GFP-hAECs were then mixed at cell ratios from 1:10 to 10:1 and centrifuged again at 200 × g for 5 minutes at room temperature. The cell pellet was resuspended in complete vascular cell media and cells were then seeded at a density of 187,500 total cells/cm^2^, based on the optimised works of Evensen *et al* [26], on Corning® 96-well Clear Flat Bottom Polystyrene TC-treated Microplates (3598).

#### 2.2.3. 3D vasculogenesis device culture

*Reagent preparation:* Fibrinogen type 1-S from bovine plasma (F8630) stock solution was prepared as per manufacturer instructions at 10 mg/mL in DPBS before being sterilised using a 0.22 µm syringe filter. Thrombin from bovine plasma (T7513) stock solution was prepared at 100 U/mL in 0.1% (w/v) bovine serum albumin solution and similarly sterilised. Aprotinin (A1153) was dissolved in DPBS to give a stock solution of 15 U/mL and sterile filtered.

*Device seeding:* Fibrinogen stock solution was diluted to 6 mg/mL in DPBS. Suspensions of GFP-hAECs and hAD-MSCs were prepared in respective cell maintenance medias and mixed at ratios from 5:1 to 1:2. Cell mixtures were centrifuged at 200 × g for 5 minutes and resuspended at 26 × 10^6^ cells/mL using a solution of 2 U/mL thrombin and 0.3 U/mL aprotinin in complete vascular cell media. 10 µL of cell suspension was quickly mixed with 10 µL of 6 mg/mL fibrinogen, then 10 µL of this mixture (13 × 10^6^ cells/mL, 3 mg/ml fibrinogen, 1 U/mL thrombin, 0.15 U/mL aprotinin) was pipetted directly into the cell seeding port of the microfluidic device. Final total cell density and hydrogel formulation was based off the works of Andrée *et al* and Wan *et al* [27, 28]. The cell-seeded device was then placed in a humidified incubator for 15 minutes at 37°C in 5% (v/v) CO_2_ in air to facilitate fibrin hydrogel formation. Media channels were then loaded with complete vascular cell media and reservoirs filled with approximately 75 µL complete vascular cell media each. Devices were finally placed in a humidified incubator at 37°C in 5% (v/v) CO_2_ in air with complete media exchanges every 2–3 days for up to 7 days.

#### 2.2.4. Time-lapse imaging

Time-lapse microscopy of both 2D and 3D vasculogenesis was conducted using a Etaluma LS720 in-incubator fluorescence microscope with automated stage control running Lumaview 720/600- Series software. Images were acquired with minimum LED intensity to minimise phototoxicity and minimum photomultiplier gain to minimise image signal-to-noise. Acquisition was conducted across multiple wells of a 96-well plate or the culture compartment of microfluidic devices using 4X and 10X objectives, respectively. Multipoint time-lapse protocols were conducted with imaging intervals of 20–30 minutes over the course of 7 days culture, capturing both FITC (excitation: 473–491 nm, emission: 502–561 nm) and brightfield channels.

#### 2.2.5. 3D vasculogenesis-adipogenesis gradient co-culture

Cell-laden fibrin hydrogels were prepared in gradient microfluidic devices as per section 2.2.3. From days 0 to 3 of culture, one side of the device was loaded with complete vascular cell media and the other with complete MSC media, with full media exchanges on day 1. On day 3, complete vascular cell media was fully exchanged for fresh complete vascular cell media and complete MSC media with a half media exchange for adipocyte initiation media (PCS-500-050). This was replicated on day 5. On day 7, complete vascular cell media was fully exchanged for fresh complete vascular cell media and adipocyte initiation media, with a half media exchange for adipocyte maturation media (PCS-500-050). This was repeated every 2–3 days to day 17. To induce vascularised adipose tissue formation across the entire length of the device, the culture gradient was then inverted on day 17. That is, adipocyte maturation media was fully exchanged for fresh complete vascular cell media and complete vascular cell media with a half media exchange for adipocyte initiation media (PCS-500-050). The process from day 3 to 17 was similarly repeated up to day 31 with this inverted configuration, a feeding strategy which is depicted in **figure 3.A**. Cultures were additionally supplemented with 1 µM spermidine (05292) or 50 nM hydroxyl(oxo)vanadium 3-hydroxypiridine-2-carboxylic acid (VO-OHpic, 10009965) for their 31-day duration.

### 2.3. Cell fixation, immunofluorescence staining and imaging

Cultures were fixed at room temperature with 3.7% (w/v) paraformaldehyde (15710) for 10 minutes then washed with DPBS three times for 5 minutes each. Fixed cultures were then permeabilized with 0.1% (w/v) Triton X-100 solution in DPBS for 10 minutes followed by three 5- minute washes with DPBS, then incubated in 10% (v/v) normal goat serum (NGS, 50062Z) in DPBS blocking solution at 4°C overnight. Cultures were then incubated in anti-CD31 mouse IgG2a (MA3100), anti-aSMA mouse IgG1 (MA5-15806), anti-CD90 rabbit IgG (MA5-32559), anti-PPARG rabbit IgG (MA5-14889), anti-Ki67 rat IgG2a (14-5698-82), and respective isotype control primary antibodies (31235, 14-4724-82 and 14-4724-82), diluted as per Table S1 in 10% (v/v) NGS, for 1 hour at room temperature, followed by three 5-minute washes with DPBS. Secondary antibodies anti-rabbit IgG AlexaFluor488 (A-11008), anti-rat IgG AlexaFluor555 (A-21434), anti-mouse IgG2a AlexaFluor555 (A-21137), anti-rabbit IgG AlexaFluor647 (A-21245), anti-mouse IgG2a AlexaFluor647 (A-21241) and anti-mouse IgG1 AlexaFluor647 (A-21240) were diluted to 4 μg/mL in 10% (v/v) NGS, or actin green solution (R37110) or actin red solution (R37112) diluted 2 drops/mL in DPBS, was incubated for 1 hour at room temperature, followed by another three 5- minute washes with DPBS. 4′,6-diamidino-2-phenylindole, dihydrochloride (DAPI) stock solution (R37606) diluted 2 drops/ml in DPBS was applied to cultures and incubated for 30 minutes at room temperature, followed by another three 5-minute washes with DPBS. For lipid staining, HCS LipidTox red neutral lipid stain (H34476) stock solution was diluted 1:200 in DPBS and applied to cultures overnight at 4°C. Images were obtained using a Nikon Ti-2 inverted fluorescence microscope or a Leica SP8 confocal laser scanning microscope. Confocal microscopy data was analysed using Imaris 10.1.0 software.

### 2.4. Culture metabolic activity assay

Total culture metabolic activity was determined via PrestoBlue™ HS Cell Viability Reagent (P50200) as per manufacturers protocol. Briefly, a 1X working solution of PrestoBlue reagent was prepared by dilution of 10X stock solution in fresh room temperature culture media. Spent media was aspirated from cultures and replaced with PrestoBlue working solution and incubated for 30 minutes at 37°C in 5% (v/v) CO_2_ in air. Following incubation, 90 µL samples of working solution was placed in Nunc™ F96 MicroWell™ Black Polystyrene Plates (237105) and fluorescence intensity determined at an excitation wavelength of 560 nm and emission wavelength of 590 nm using an Ensight^TM^ Multimode Micro Plate Reader (Perkin Elmer).

### 2.5. Quantitative image analysis

#### 2.5.1. Vessel network morphology

Fluorescence images of GFP-hAECs were obtained as described in section 2.2.4 using a 4X objective with consistent illumination intensity, photomultiplier gain and exposure time. Images were captured with maximum vessel features in the plane of focus. Quantification of 2D and 3D vascular network morphology (including number of end points, number of junctions, total vessel length and total vessel area) was conducted using AngioTool version 0.6a [29], a lightweight open-source software for quantitative analysis of vessel networks. Vessel network analysis was conducted at a user defined intensity scales which maximised accuracy of vessel network segmentation. Single cells and other fluorescent non-vessel particles were excluded from the analysis using the software’s ‘remove small particles’ function and small holes and dark spots within vasculature were filled using the ‘fill holes’ function.

#### 2.5.2. Vessel diameter quantification

Vessel diameters were measured in Fiji [30] by manually drawing a line the thickness of a vessel approximately equidistant from two junctions. Pixel length was then converted into a distance measurement using a known scale reference.

#### 2.5.3. Lipid quantification

Colour or monochromatic images of 3D vasculogenesis-adipogenesis gradient cultures were obtained at days 17 and 31 of culture using an Olympus CKX 53 or Nikon Ti-2 microscope. Images were captured at consistent illumination and gain values using the autoexposure camera function. Images were imported into ImageJ Fiji software and converted to 8-bit format. Image threshold was adjusted to highlight dark lipid droplet-like structures with spherical morphology in non-vessel control cultures. These threshold parameters were used to analyse all other images. Regions of interest within the culture images were manually selected and percentage area of image within the threshold values determined.

### 2.6. Growth factor distribution modelling

Idealised growth factor distributions were simulated within the fibrin hydrogel culture compartment of the gradient microfluidic device (**figure 3**). Following Altan-Bonnet & Mukherjee [31], growth factors were assumed to degrade throughout culture *via* half-life decay and transport from microfluidic channels into the hydrogel culture compartment *via* diffusion. In this simple model, we assume growth factor media supplementation and half-life decay to be significantly greater than other kinetics, such as growth factor secretion and cell or matrix sequestering, as found in our prior work modelling single-cell cytokine release and binding [32]. The composition of adipocyte initiation and maturation media is proprietary and therefore was approximated from the following source [33]. Therefore, 2D growth factor transport and decay over time within the gradient microfluidic device was simply modelled as:

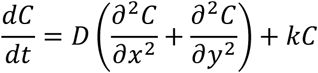

Where, growth factor concentration *C* is modelled as a partial differential equation of dimensions *x*, *y* and *t*. Here, *D* represents a growth factor’s diffusivity constant (µm^2^/s) and half-life decay rate *k* is equal to *ln(0.5/t_1/2_)* with *t_1/2_* (s) being the time for the growth factor to decay to half its original concentration. It is important to note this simulation also applies the same half-life decay to the microchannel boundary concentration conditions, as media is only replenished every 2–3 days.

The microchip’s 3D fibrin hydrogel cell culture chamber is 1.3 mm wide and 7.9 mm long, where four media channels interface with the culture chamber at its top-left, top-right, bottom-left, and bottom-right corners for 2.6 mm of length, each. **Table 1** lists major growth factors’ initial conditions, diffusivity, half-life, half maximal effective concentration (EC50) from these experiments. It is important to note that these parameters were derived from 2D culture conditions and growth factor transport, decay and effect would likely change in hydrogels [34]. Further, diffusion or half-life kinetics for adipogenic factor IBMX (3-isobutyl-1-methylxanthine) were unable to be identified and modelled.

**Table 1:**
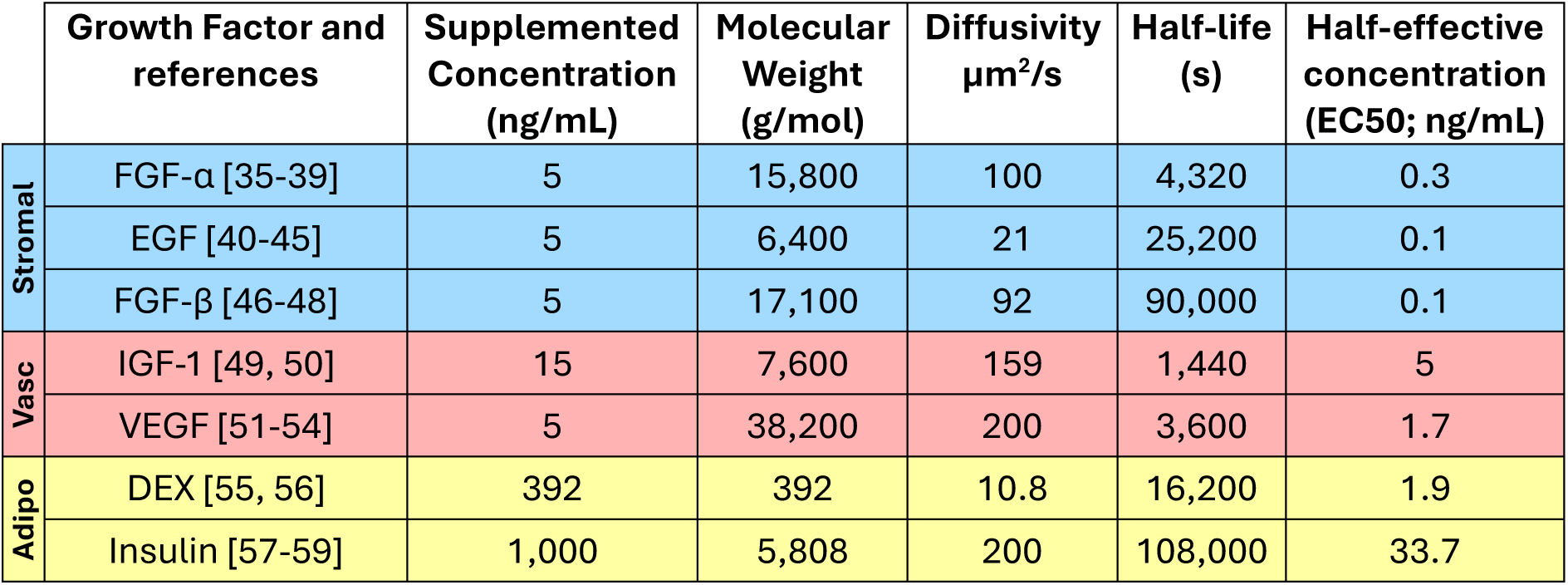
Growth factor distribution modelling parameters. Blue, red, and yellow rows represent critical stromal, vascular, and adipogenic growth factors, respectively. It’s important to note that EGF and FGF-β are included in both vascular cell and MSC medias. IGF-1: insulin-like growth factor 1, VEGF: vascular endothelial growth factor, EGF: epidermal growth factor, FGF-β: fibroblast growth factor-basic, FGF-α: fibroblast growth factor-acidic, DEX: dexamethasone.

| | Growth Factor and references | Supplemented Concentration (ng/mL) | Molecular Weight (g/mol) | Diffusivity $\mu\text{m}^2/\text{s}$ | Half-life (s) | Half-effective concentration (EC50; ng/mL) |
| --- | --- | --- | --- | --- | --- | --- |
| Stromal | FGF-α [35-39] | 5 | 15,800 | 100 | 4,320 | 0.3 |
|  | EGF [40-45] | 5 | 6,400 | 21 | 25,200 | 0.1 |
| | FGF-β [46-48] | 5 | 17,100 | 92 | 90,000 | 0.1 |
| Vasc | IGF-1 [49, 50] | 15 | 7,600 | 159 | 1,440 | 5 |
|  | VEGF [51-54] | 5 | 38,200 | 200 | 3,600 | 1.7 |
| Adipo | DEX [55, 56] | 392 | 392 | 10.8 | 16,200 | 1.9 |
|  | Insulin [57-59] | 1,000 | 5,808 | 200 | 108,000 | 33.7 |

The distribution of each growth factor was individually simulated across the microfluidic cell culture compartment in figure S17. FGF-α, IGF-1, and DEX were respectively identified to be the limiting growth factors for stromal, vasculogenic, or adipogenic media distribution. That is, these growth factors were predicted to distribute the shortest distance across microfluidic chip culture, while still maintaining bioactive concentrations. A short growth factor distribution could be due to smaller diffusivity, shorter half-life, or lower supplemented concentration relative to higher half-effective concentrations (EC50) [31]. Detailed simulations of the limiting growth factors (FGF-α, IGF-1, DEX) were performed in **figure 3**. The Python simulation code is provided in the supplementary information for an exemplary pair of counter-current growth factors, where its approach in implementing a 2D Laplacian for growth factor diffusion follows the methodology of Rossant [60].

### 2.7. Statistical analysis

Unless otherwise stated in the figure caption or text, all experiments included three biological replicates, each with three technical replicates. Results are reported as an average of biological replicates ± standard deviation. Statistical analysis and multiple comparison tests are detailed in figure captions. P value output style is P > 0.05 (ns), P ≤ 0.05 (*), P ≤ 0.01 (**), P ≤ 0.001 (***) and P ≤ 0.0001 (****).

## 3. Results

### 3.1. hAECs and hAD-MSCs spontaneously form robust vessel-like networks in optimised 2D culture conditions

Immunocytochemical detection confirmed the expression of the phenotypical markers CD31 and CD90 in GFP-hAEC and hAD-MSC monocultures, respectively. CD31 was observed to stain plasma membrane and cell nucleus of sub-confluent GFP-hAEC cultures (figure S1) and CD90 similarly in sub-confluent hAD-MSC cultures (figure S2). hAD-MCSs were highly proliferative, with 86.27% of hAD-MSCs expressing Ki67 (figure S2.B).

The ability of these two cell types to spontaneously self-assemble into vascular networks was investigated by simultaneously seeding GFP-hAEC and hAD-MSCs into uncoated tissue culture polystyrene at a density of 187,500 total cells/cm^2^ with GFP-hAEC:hAD-MSC ratios of 2:1, 1:1, 1:2, 1:5 and 1:10 in complete vascular cell media for 7-days (**figure 1.A, video S1**). Cultures with seeding ratios of 2:1 and 1:1 were observed to commence vessel network formation, however, detached from plates prematurely as early as approximately 22- and 82-hours post-seeding, respectively. Cultures with seeding ratios 1:2, 1:5 and 1:10 formed robust vessel networks which sustained attachment throughout the entire 7-day culture period (**figure 1.A.iii-v**). The number of discontinuities in vessel networks after 7 days culture was observed to increase with decreasing proportion of GFP-hAECs in the cell seeding mixture (**figure 1.A.v**).

**Figure 1.**
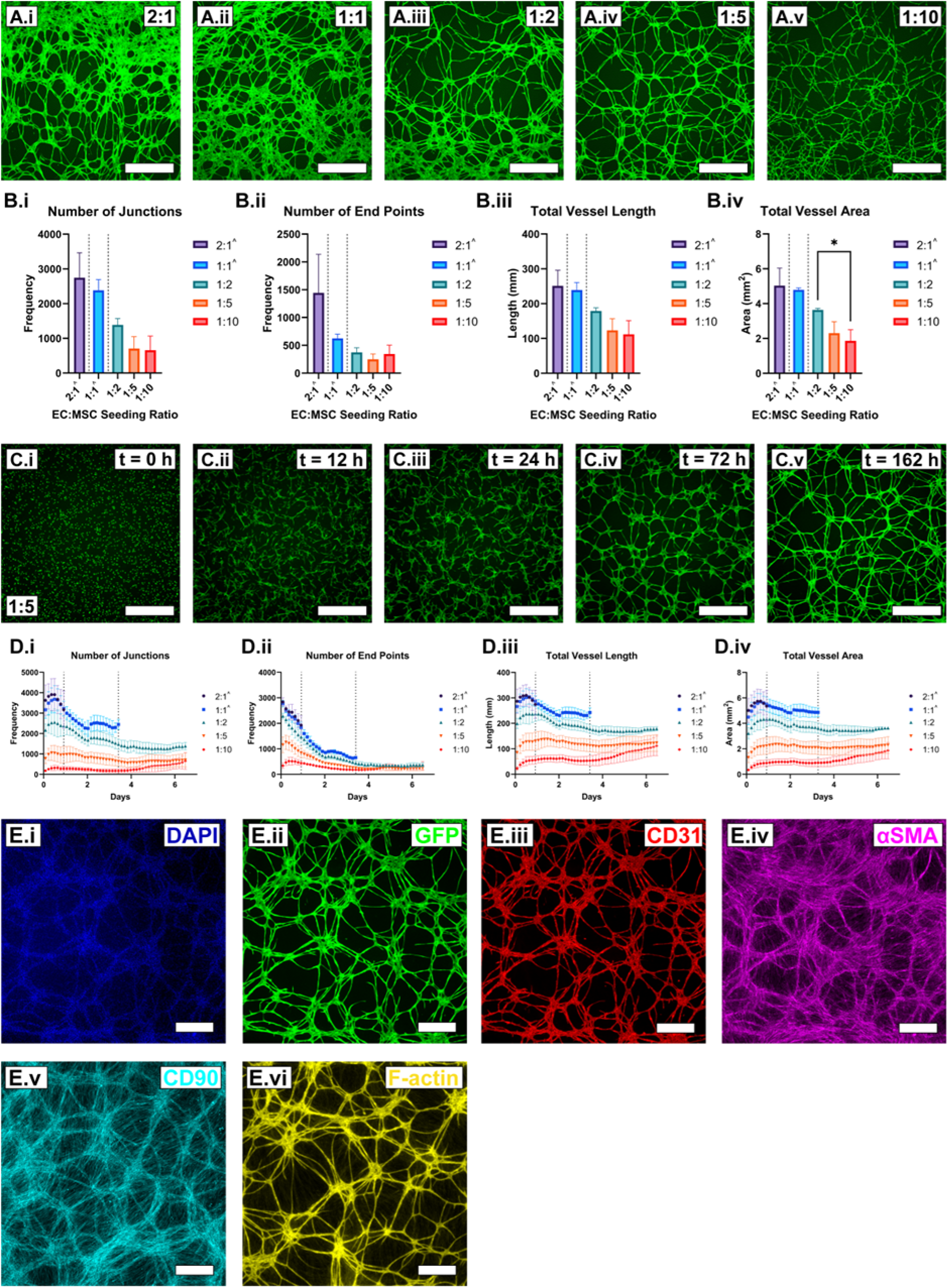
GFP-hAECs and hAD-MSCs spontaneously form robust and phenotypical 2D microvascular networks with a morphological dependence on cell seeding ratio. A. Fluorescence images of GFP-hAEC microvascular networks formed after 7-days culture in complete vascular cell media, with GFP-hAEC:hAD-MSC seeding ratios of 2:1 (i), 1:1 (ii), 1:2 (iii), 1:5 (iv) and 1:10 (v, scale bars = 1 mm). B. Quantitative analysis of vessel network morphology at day 7, detailing the number of network junctions (i), vessel end points (ii), total vessel length (iii) and total vessel area (iv) within a field of view 3.67 × 3.67 mm (N = 3 biological replicas, mean ± standard deviation). Results were compared via an ordinary one-way ANOVA followed by a Tukey multiple comparisons test. C. Time lapse imaging of GFP-hAECs in 7-day vasculogenesis co-culture with a seeding ratio of 1:5 (scale bars = 1 mm). D. Quantitative analysis of every tenth time-lapse image (47 of 478 total images) over the entire 7-day culture period with graphs depicting the number of network junctions (i), vessel end points (ii), total vessel length (iii) and total vessel area (iv) within a field of view 3.67 × 3.67 mm (N = 3 biological replicas, mean ± standard deviation). E. Immunocytochemical staining panel of a 1:5 seeding ratio sample after 7-days culture detailing the nuclear counter stain DAPI (i), endogenous GFP expression (ii), and detection of CD31 (iii), αSMA (iv), CD90 (v, false coloured cyan) and F-actin (vi, false coloured yellow, scale bars = 500 µm). ˄At times, the 2:1 and 1:1 seeding ratio conditions detached from culture surfaces. The minimum time directly before which a culture commenced detachment (2:1. 22-hours; 1:1, 82-hours) was used for the analysis of that condition and therefore was not statistically compared to the cultures which remained attached for the culture duration. Depicted as dotted line in 1.B and D. EC: endothelial cell, MSC: mesenchymal stem cell, DAPI: 4′,6-diamidino-2-phenylindole, GFP: green fluorescent protein, CD31: cluster of differentiation 31, αSMA: alpha smooth muscle actin, CD90: cluster of differentiation 90, F-actin: filamentous actin.

At culture termination, the number of end points, number of junctions, total vessel length and total vessel area generally decreased with smaller GFP-hAEC:hAD-MSC seeding ratios, with the exception of the 1:10 ratio, which displayed a greater average number of end points than the 1:5 seeding ratio (**figure 1.B.ii**). The time points used for quantification of the 2:1 and 1:1 ratios, which prematurely detached before the 7-day culture endpoint, were defined as the earliest detachment time of any replicate of that condition (2:1, t = 0.92 days; 1:1, t = 3.42 days).

Time-lapse microscopy supplemented with image analysis was used to quantitatively examine the number of vessel network junctions and endpoints as well as the total vessel area and length over a 7-day culture period (**figure 1.C-D, figure S3**). All four of these metrics demonstrated a rapid increase in magnitude, reaching a maximum at approximately 12 hours post seeding. After which, these properties began decreasing as individual endothelial cells joined together to form microvascular networks, stabilising after approximately 4 days in culture (**figure 1.C**). Post 4-days culture, the 1:10 seeding ratio culture condition exhibited a further gradual increase in all four measured vessel metrics, which coincided with an observed increase in angiogenic-like sprouting from established GFP-expressing vessel-like structures from day 4 up to day 7 (figure S4). Angiogenic-like sprouting was observed from existing vessel-like structures throughout culture (figure S5.A) and vessel-like structures were also observed to merge end-to-end (figure S5.B). No observable signs of abnormal cell function resulting from phototoxicity during timelapse imaging for the length of the 7-day culture period were observed, when compared to non-imaged cultures.

At day 7, 2D vasculogenesis cultures were fixed and immunocytochemically stained for the detection of phenotypical human vascular network markers (**figure 1.E, figure S6**). DAPI nuclear staining was observed in nuclei-like structures through the culture plate and GFP was detected exclusively in vessel-like structures (**figure 1.E.i-ii**). Staining for the expression of CD31 was observed exclusively on vessel-like structures, with staining most intense at cell-cell junctions (**figure 1.E.iii**). CD90 staining was observed in cells lining the bottom of the plate and cells lining vessel-like structures (**figure 1.E.iv**). Staining for the expression of αSMA was observed strongly in cells lining vessel-like structures and less so in cells attached the bottom of the culture plate (**figure 1.E.v**). Chemical staining for filamentous actin was observed in vessel-like structures and comparatively less in extravascular space (**figure 1.E.vi**), similar to other 2D co-cultures [61]. At higher magnification, cells positive for αSMA were observed to line and bridge GFP-positive vessel-like structures (figure S7).

Vasculogenic co-cultures require hAD-MSCs to be cultured in media different to manufacturer specifications. To characterise how different media factors effect GFP-hAECs and hAD-MSCs, monocultures were examined for metabolic activity in their respective complete and basal culture medias, as well as hAD-MSC monocultures in complete vascular cell media (figure S8). GFP-hAEC and hAD-MSC monocultures in their respective complete medias demonstrated an increase in culture metabolic activity up to day 5 which was maintained to day 7. Both GFP-hAEC and hAD-MSC monocultures in respective basal medias displayed a reduction in culture metabolic activity across the entire 7-day culture period. hAD-MSC monoculture metabolic activity was found to increase in complete vascular cell media compared to complete MSC media. The vasculogenic potential of GFP-hAECs was then investigated in the absence of complete vascular cell media supplements and also the absence of hAD-MSCs, and compared to a 1:5 GFP-hAEC:hAD-MSC control in complete vascular cell media (figure S9). In the absence of hAD-MSCs (cultured in complete vascular cell media), GFP-hAECs formed a confluent monolayer with little evidence of vessel-like morphology (figure S9.A). In the absence of vascular cell media supplements, GFP-hAEC were observed not to attach to the tissue culture polystyrene culture surface (figure S9.B). In control conditions, vessel network formation proceeded typically (figure S9.C).

### 3.2. Microfluidic culture conditions support spontaneous formation of 3D fibrin hydrogel-embedded GFP-hAECs and hAD-MSCs into robust and patent vessel-like networks

The ability of GFP-hAECs and hAD-MSCs to spontaneously form robust vessel networks embedded in 3D fibrin hydrogels was investigated using GFP-hAEC:hAD-MSC seeding ratios of 5:1, 2:1, 1:1 and 1:2 (**figure 2.A**) within previously validated microfluidic device configurations [28, 61, 62] (figure S10). Total vessel length, number of junctions and number of endpoints were all found to increase with decreasing GFP-hAEC:hAD-MSC seeding ratio (**figure 2.B.i-ii, iv**), however, total vessel areas were found to decrease proportionally with decreasing GFP-hAEC:hAD-MSC seeding ratio (**figure 2.B.iii**). Vessel diameter was also qualitatively observed to decrease with decreasing GFP-hAEC:hAD-MSC seeding ratio (**figure 2.A**). Vessel morphology was observed to be consistent across the entire length of the culture compartment (figure S11).

**Figure 2.**
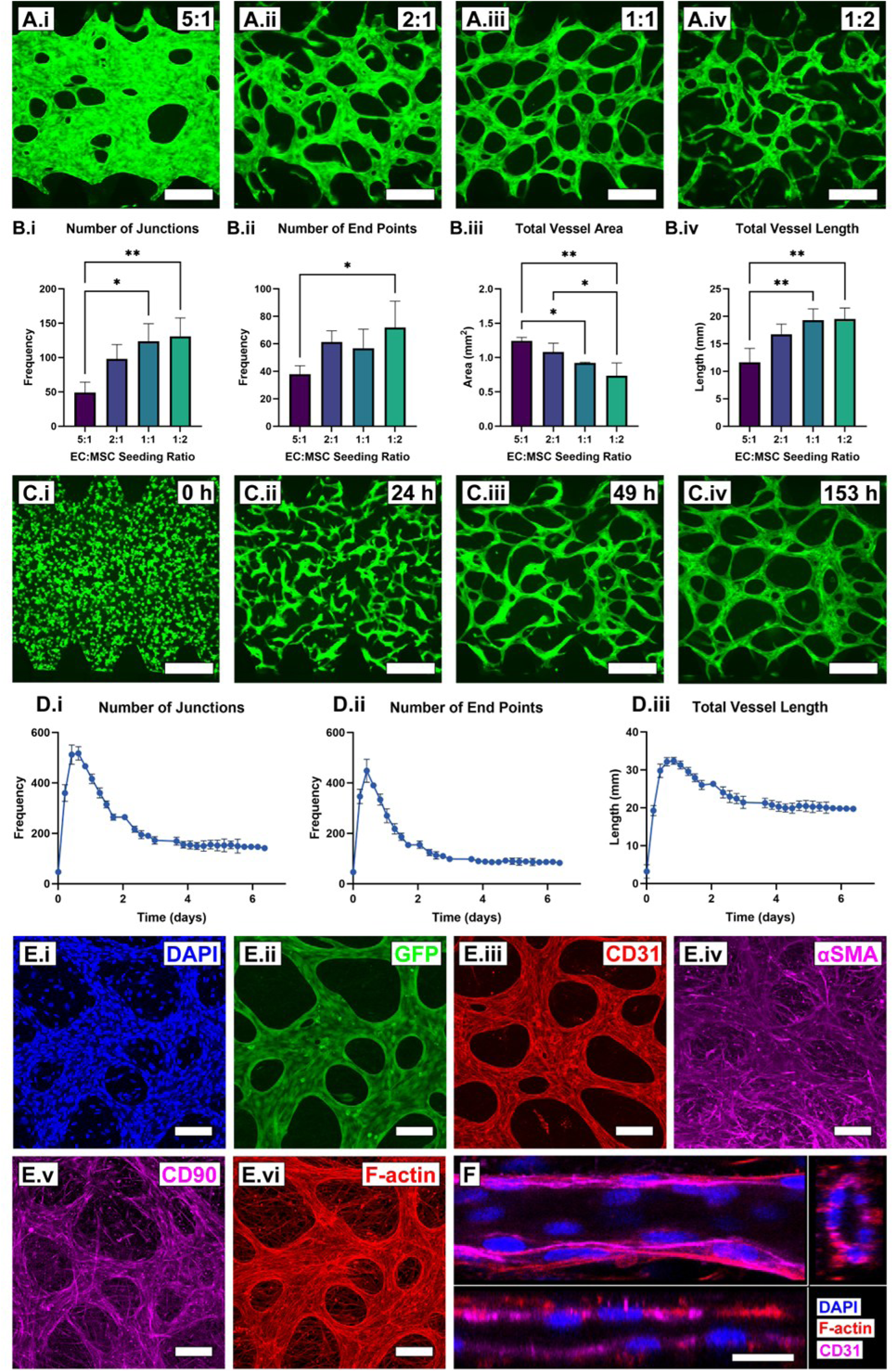
GFP-hAECs and hAD-MSCs spontaneously form robust and phenotypical 3D hydrogel-embedded microvascular networks with a morphological dependence on cell seeding ratio. **A.** Fluorescence images of GFP-hAECs as microvascular networks formed after 7-days fibrin hydrogel-embedded microfluidic device culture in complete vascular cell media with GFP-hAEC:hAD-MSC seeding ratios of 5:1 (i), 2:1 (ii), 1:1 (iii) and 1:2 (iv, scale bars = 300 µm). B. Quantitative analysis of vessel network morphology detailing the number of network junctions (i), vessel end points (ii), total vessel length (iii) and total vessel area (iv) within a 1.33 × 1.33 mm field of view (N = 3 biological replicas, mean ± standard deviation). Results were compared via an ordinary one-way ANOVA followed by a Tukey multiple comparisons test. C. Time lapse imaging of GFP-hAECs in 7-day vasculogenesis co-culture with a seeding ratio of 1:1 after 0 (i), 24 (ii), 49 (iii) and 153 (iv) post-seeding (scale bars = 300 µm). D. Quantitative analysis of every tenth time-lapse image (28 of 274 images total) of the entire 7-day culture period, with graphs depicting the number of network junctions (i), vessel end points (ii) and total vessel length (iii) within a 1.33 × 1.33 mm field of view (mean ± standard deviation). E. Immunocytochemical characterisation of a 1:1 seeding ratio sample after 7-days culture, detailing the nuclear counter stain DAPI (i), endogenous GFP expression (ii), and detection of CD31 (iii), αSMA (iv), CD90 (v) and F-actin (vi, scale bars = 100 µm). F. DAPI nuclear counter stain, CD31 and F-actin sample cross sections demonstrating vessel patency (scale bar = 20 µm). EC: endothelial cell, MSC: mesenchymal stem cell, DAPI: 4′,6-diamidino-2-phenylindole, GFP: green fluorescent protein, CD31: cluster of differentiation 31, αSMA: alpha smooth muscle actin, CD90: cluster of differentiation 90, F-actin: filamentous actin.

When analysed via time-lapse microscopy, with imaging intervals every 30 minutes over a 7-day period (**figure 2.C, video S2-3**), the 1:1 seeding ratio condition demonstrated an increase in number of junctions, number of end points, and total vessel length up to approximately 12 hours post-seeding, after which, these values began to decrease and plateau after approximately 4- days and remained constant up to day 7 (**figure 2.D**). Total vessel area could not be accurately quantified due to discrepancies and changes in GFP intensities across the surfaces of 3D vessel networks over time.

After 7 days, 3D vasculogenesis cultures were fixed and immunocytochemically stained for the detection of phenotypical vascular network markers (**figure 2.E**). Confocal microscopy revealed the detection of CD31 expression in vessel-like structures, most intensely staining the cell-cell junctions (**figure 2.E.iii**). CD31 detection was also found in some spherical cells observed inside vessel-like structures, potentially being shed endothelial cells or those preparing to divide (figure S12). αSMA and CD90 detection was observed similarly in cells both lining vessel-like structures and occupying extra-vascular space (**figure 2.E.iv-v**). F-actin staining revealed the detection of filament-like structures, distributed both in the parenchymal space between and on vessel-like networks (**figure 2.E.vi**). Confocal microscope imaging and 3D reconstruction of vessel-like structures revealed an outer lumen-like structure stained positive for the detection of CD31 and actin and a vacant inner lumen absent of staining (**figure 2.F**). This morphology was found to be consistent throughout the vascular-like network and the intra-luminal space was determined to be 9.76 ± 2.11 µm on average in the z-direction. Cell nuclei were observed to distribute throughout the entire space of a 582 × 582 × 35.6 µm imaging volume encompassing the microvascular network (figure S13.A). Within this imaged volume, 25.8% of all segmented nuclei (1514 total nuclei) were found to be non-overlapping (overlap score = 0) with the microvascular network structure, indicating non-mural parenchymal cell fraction (figure S13.B).

To determine GFP-hAEC dependency on supplemented media versus hAD-MSC paracrine-supported growth, the 3D vasculogenic potential of GFP-hAECs was assessed in the absence of hAD-MSCs, the absence of complete vascular cell media supplements and compared to a 1:1 seeding ratio control in complete vascular media (figure S14). 3D cultures of GFP-hAECs (6.5 × 10^6^ cells/mL) in the absence of hAD-MSCs still formed vessel-like networks after 7-days culture, however, networks displayed discontinuities and GFP-expressing cell borders were rougher in morphology than typically observed in the presence of hAD-MSCs (figure S14.A). When cultured in vascular cell basal media, at a 1:1 seeding ratio, GFP-hAECs still formed continuous vessel-like networks with smooth GFP-expressing cell borders (figure S14.B). Vessel-like networks formed in vascular cell basal media exhibited an average diameter 19.1 ± 3.72 µm (figure 14.B), less than control cultures 34.22 ± 1.79 µm (figure S14.C).

An alternative seeding method was explored to aid the formation of vessel openings between microfluidic device pillars. A suspension of 13 × 10^6^ GFP-hAECs/mL was prepared in a hydrogel precursor solution as described in section 2.2.3, pipetted into the device and then immediately aspirated (figure S15.A.i). A second hydrogel precursor solution was then loaded with 13 × 10^6^ total cells/mL with a GFP-hAEC:hAD-MSC ratio of 1:1 (figure S15.ii-iii). This resulted in an increased vessel width in between pillars (figure S15.B) compared to control cultures (figure S15.C), however, did not support the perfusion of a fluorescently-labelled dextran solution throughout vessel networks (Dextran Cascade Blue^TM^, 10,000 MW, D1976; data not shown).

### 3.3. Microvascular network co-cultures enhance hAD-MSC adipogenic differentiation and subsequent lipid production

The adipogenic potential of hAD-MSCs was initially validated in 2D well plate formats according to manufacturer’s protocol for 17 days total. After 17 days culture, spherical lipid-like droplets were visible via brightfield microscopy clustered together and surrounding some cells (figure S16.A.iv). Lipid morphology was confirmed via staining with the fluorescent lipid-sensitive dye, LipidTOX (figure S.16.A.i-iii). Adipogenesis cultures also demonstrated the detection of preadipocyte marker PPARG via immunocytochemical staining at day 17 (figure S.16.B-C).

Two-channel microfluidic devices (figure S10) were modified to include four channels to support nutrient gradient formation across the device (**figure 3.B**). Numerical modelling of growth factor and small molecule diffusion and decay validated distribution across the length of the device (**figure 3.C**). The diffusion-decay of all eight major growth factors in complete MSC media, complete vascular cell media, and adipogenic initiation/maturation culture media were simulated and IGF-1, FGF-α, and DEX were found to be the most distribution-limited factors for each culture media, respectively (figure S17).

**Figure 3.**
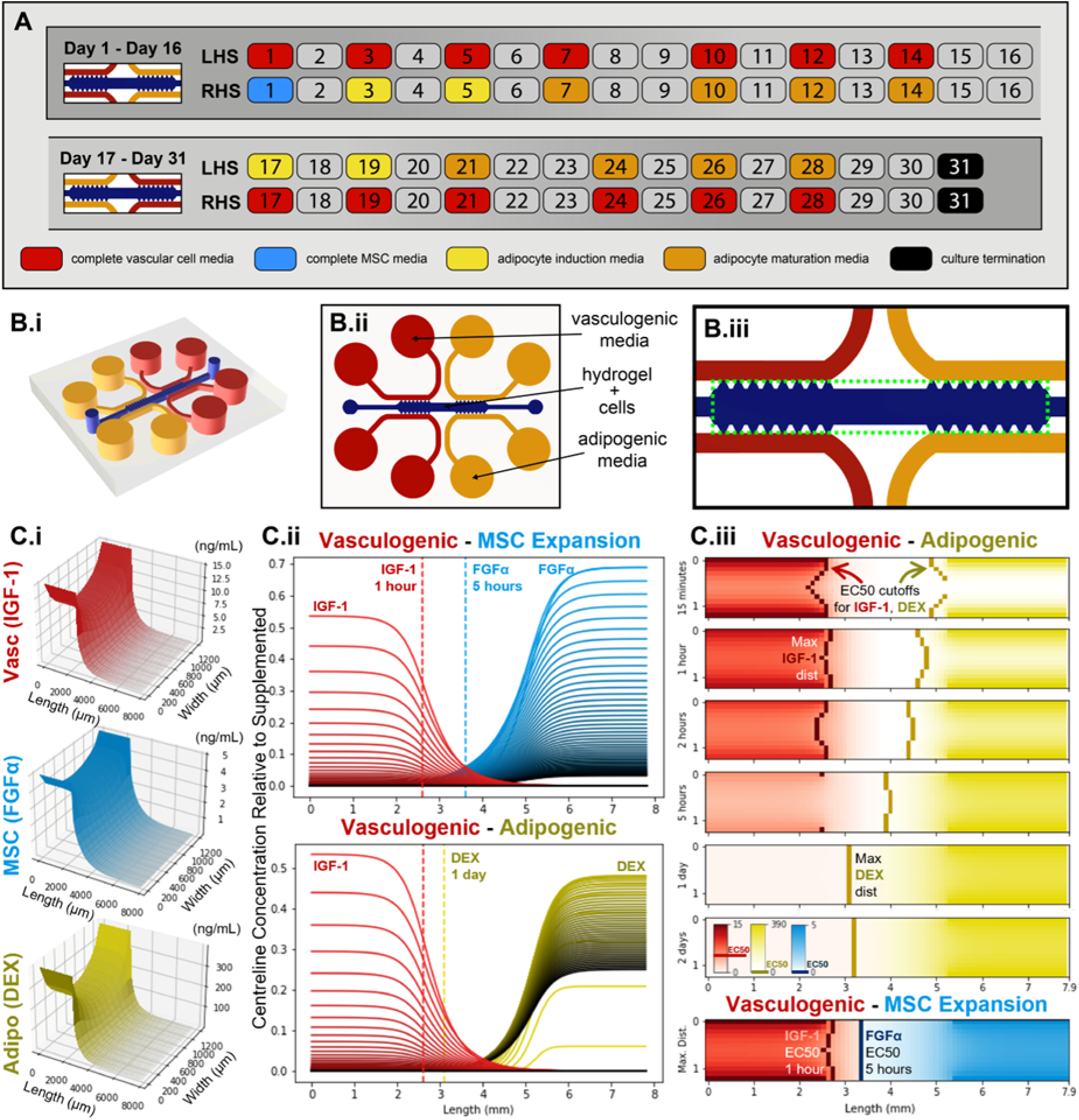
Simulated 3D hydrogel microfluidic co-culture design to generate counter-current gradients of vasculogenic and adipogenic factors. **A.** Gradient device culture typical feeding strategy depicting each day of a complete 31-day culture. Both adipocyte induction (light yellow) and maturation media (dark yellow) represent half media exchanges as per section 2.2.5. **B**. The hydrogel culture compartment (dark blue) is fed either end by adipogenic (yellow) and vasculogenic cell culture media (red; i-ii). The culture compartment area (iii, green dotted square) is modelled for molecular diffusion. **C**. The most distribution-limited vascular (Vasc, red), stromal (MSC, light blue), adipogenic (Adipo, yellow) growth factors were found to be IGF-1, FGF-α, and DEX. The maximum simulated concentration (in ng/mL) of these factors for each mm^2^ of culture compartment area (13 × 79 mm) over 2 days (i). Growth factor concentrations across the culture compartment length at mid-width, relative to supplemented concentrations, where line colour turns darker for each hour of culture up to 2 days (ii). Growth factor concentration heatmaps over culture compartment area and time for the vasculogenic-adipogenic condition, then maximum distribution of the vasculogenic-MSC expansion condition at bottom (iii). Vertical lines in C.ii and C.iii represent the area where the simulated growth factor concentration is at EC50. Detailed simulations for each growth factor can be found in figure S17, and representative code used to generate figure 3.C can be found in the supplementary materials. LHS: left hand side, RHS: right hand side, IGF-1: insulin-like growth factor 1, FGF-α: fibroblast growth factor acidic, DEX: dexamethasone.

The fraction of the microfluidic chip’s culture area predicted to receive a bioactive concentration of a growth factor (above its EC50) was used to compare growth factor distributions (**figure 3.B.ii-iii; figure S17**). Simulations predicted that vasculogenic factors distributed the shortest length into the culture compartment; a maximum distribution of 2.6 mm and 3.1 mm for IGF-1 or VEGF, respectively. In contrast, stromal growth factors FGF-α, EGF, or FGF-β exhibited distributions of 4.2 mm, 5.4 mm and 7.9 mm (the entire microchip), respectively. Adipogenic factors distribute furthest along the chip, with dexamethasone at 4.7 mm and insulin at 7.9 mm (the entire length of the culture compartment). These predicted distributions were shorter for growth factors with a smaller supplementation-to-EC50 concentration ratios or a shorter half-life (IGF-1, VEGF, FGF-α at 24, 60, and 72 minutes). Growth factors with a short half-life experienced maximum distribution soon after media addition (IGF-1, VEGF, FGF-α at 0.5, 1, and 5 hours, respectively) and then soon after resided at sub-bioactive concentrations between media exchanges, as compared to factors with a long half-life (EGF, FGF-β, insulin at 1, 2, and 2 days, respectively).

After 17-days of gradient culture (**figure 4.A**), a distribution of spherical lipid-like droplets was qualitatively observed across the device culture compartment using colour brightfield microscopy (**figure 4.A.i, figure S18**). The appearance of lipid-like droplets was more apparent in GFP-hAEC and hAD-MSC co-cultures of all ratios (1:2, 1:1 and 2:1) when compared to the hAD-MSC monoculture condition (**figure 4.B**). After 31 days culture (following media inversion at day 17), the abundance and density of lipid-like droplet in culture continued to increase (**figure 4.A.ii-iii**). Fluorescent lipid staining and confocal microscope imaging confirmed the presence of lipid droplets throughout the culture chamber (**figure 4.C**). Quantitative analysis of colour brightfield images revealed a lipid coverage of 67.4 ± 0.07%, 62.67 ± 8.52% and 56.1 ± 15.6% in the 1:2, 1:1 and 2:1 co-culture conditions, respectively; compared to 1.86 ± 1.09% in the hAD-MSC mono-culture condition (**figure 4.D**). Additionally, immunocytochemical characterisation revealed the detection of PPARG after 17 days of culture (**figure 4.E, figure S19**).

**Figure 4.**
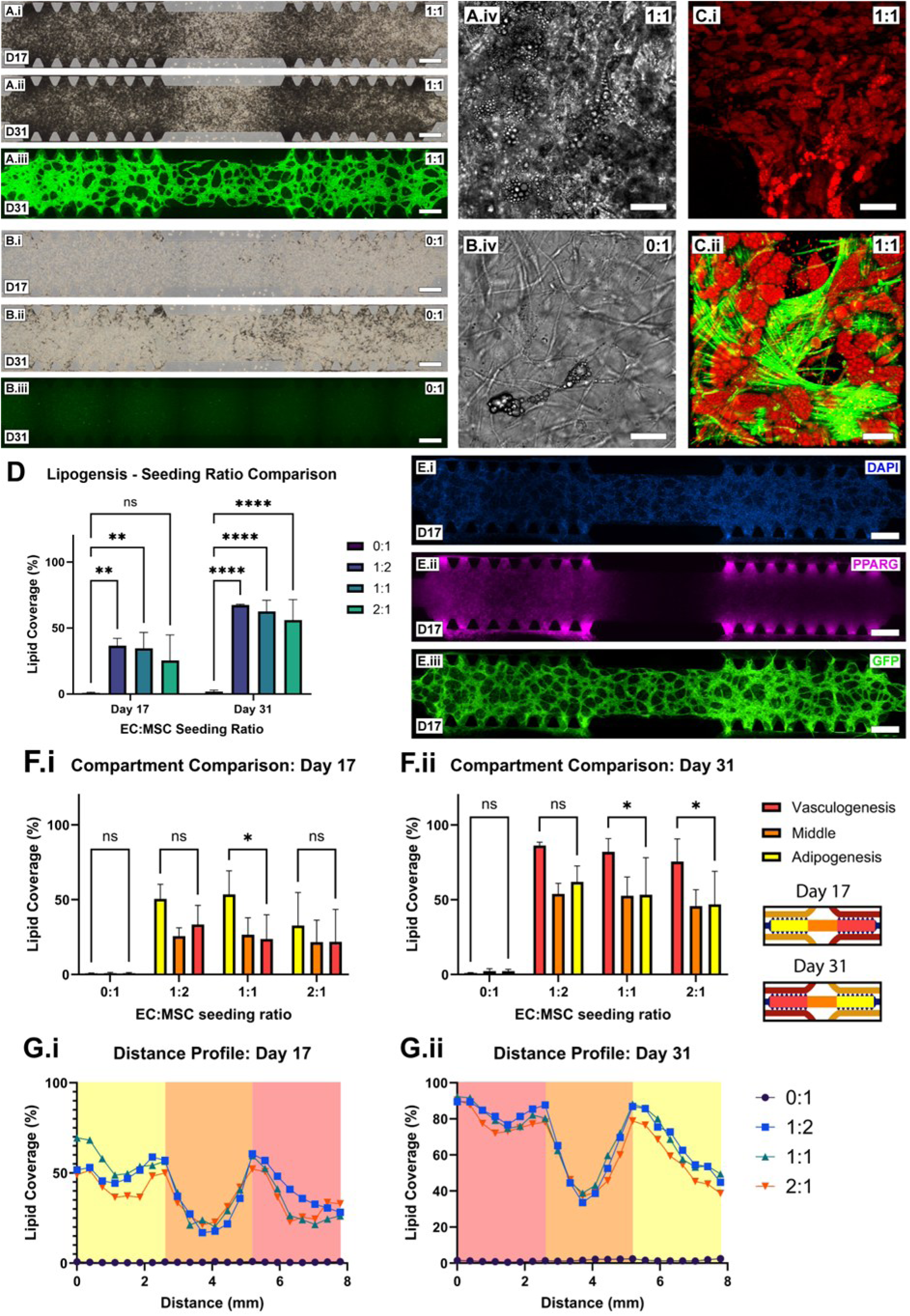
GFP-hAEC and hAD-MSC gradient co-culture supports the co-formation of microvascular networks and differentiation with vascular network-enhanced adipogenesis. **A.** Colour brightfield microscopy images of 1:1 seeding ratio microfluidic device cultures after 17-(i) and 31-days (ii) gradient culture and green fluorescence after 31 days culture (iii, scale bars = 500 µm). High magnification monochromatic brightfield image of the culture compartment (iv, scale bar = 50 µm). **B.** Colour brightfield microscopy images of 0:1 seeding ratio microfluidic device cultures after 17-(i) and 31-days (ii) gradient culture and green fluorescence after 31 days culture (iii, scale bars = 500 µm). High magnification monochromatic brightfield image of the culture compartment (iv, scale bar = 50 µm). **C.** LipidTOX staining of 1:1 seeding ratio cultures at day 31 at low magnification (i, scale bar = 50 µm) and high magnification with F-actin (green) and GFP (ii, scale bar = 30 µm). **D.** Quantitative analysis of lipid coverage across the culture compartment at days 17 and 31 comparing seeding ratios of 0:1, 1:2, 1:1, 2:1 (N = 3 biological replicas, mean ± standard deviation). Results were compared via an ordinary two-way ANOVA followed by a Tukey multiple comparisons test. **E.** Immunocytochemical detection of 1:1 co-cultures at day 17 for PPARG with images depicting the nuclear counter stain DAPI (i), PPARG (ii) and GFP-hAECs (iii, scale bars = 500 µm). **F.** Percentage lipid coverage localised to each third of the culture compartment defined as vasculogenesis, middle and adipogenesis (inset). Seeding ratios of 0:1, 1:2, 1:1 and 2:1 were investigated at days 17 and 31 of culture (N = 3 biological replicas, mean ± standard deviation). Results were compared via an ordinary one-way ANOVA followed by a Tukey multiple comparisons test. **G.** Lipid coverage was similarly quantified in 370 µm increments across the entire length of the culture compartment at day 17 and 31 of culture and plotted for seeding ratios of 0:1, 1:2, 1:1 and 2:1 (N = 3 biological replicas, mean). EC: endothelial cell, MSC: mesenchymal stem cell. EC: endothelial cell, MSC: mesenchymal stem cell, D: day, F-actin: filamentous actin, DAPI: 4′,6-diamidino-2-phenylindole, PPARG: peroxisome proliferator-activated receptor gamma, GFP: green fluorescent protein.

When quantifying the lipid coverage per culture compartment (**figure 4.F**), a significant difference between the adipogenesis and vasculogenesis sides of the device after 17 days was observable in the 1:1 co-culture condition (**figure 4.F.i**). After 31 days culture, significant differences in lipid coverage were observed in both 1:1 and 2:1 conditions and not in the 1:2 condition (**figure 4.F.ii**). There was found to be no significant difference in lipid coverage between the adipogenesis and vasculogenesis sides of the device for the 0:1 monoculture condition at either day 17 or day 31. This distribution of lipid coverage across the length of the culture chamber was found by quantitative image analysis to be relatively constant across the first third of the device, demonstrated a symmetrical dip in the middle third and decreased in the final third for all co-culture ratios at both days 17 and 31 post-seeding (**figure 4.G**). The monoculture condition was consistently distributed in lipid coverage along the length of the culture compartment at both days 17 and 31 post-seeding.

### 3.4. Interrogation of vessel-mediated lipogenesis

Modulation of lipogenesis in gradient co-cultures was investigated using know inhibitors of adipogenesis and inducers of lipolysis, VO-OHpic and spermidine [4, 63]. To ensure VO-OHpic and spermidine had no influence on vasculogenic cells, monocultures of GFP-hAECs and hAD- MSCs were tested for metabolic activity in the presence of respective growth media supplemented with 0, 5, 50 and 500 nM VO-OHpic and 0, 0.1, 1 and 10 µM spermidine. Culture metabolic activity of either cell type was not significantly influenced by any of the investigated molecule concentrations (figure S20). Both 1:1 co-cultures and 0:1 monocultures were conducted individually in the presence of 1 µM spermidine, 50 nM VO-OHpic and in their absence (**figure 5.A, figure S.21**). The 1:1 and 0:1 control condition was conducted only with two biological replicas due to a technical error. All remaining conditions encompassed three biological replicates. Captured monochromatic brightfield images of 1:1 co-cultures at day 31 demonstrated a typical dark shade of spherical droplets distributed throughout the culture compartment of control and similarly in 1 µM spermidine and 50 nM VO-OHpic conditions (**figure 5.B.i-iii**). Brightfield images of all monocultures conditions demonstrated a consistent light shade with some small and sparse dark regions (**figure 5.B.iv**). Co-cultures were determined via quantitative image analysis to demonstrate a lipid coverage of 85.9 ± 0.07%, 85.9 ± 4.23% and 87.9 ± 6.15% for control, 1 µM spermidine and 50 nM VO-OHpic conditions, respectively (**figure 5.C**). No difference was observed between the control culture condition and either of the supplemented culture conditions. Similarly, monocultures demonstrated a lipid coverage of 0.25 ± 0.28%, 1.03 ± 0.25%, and 1.25 ± 1.34% for control, 1 µM spermidine and 50 nM VO-OHpic conditions respectively and no difference was observed between the control culture condition and either of the supplemented culture conditions.

**Figure 5.**
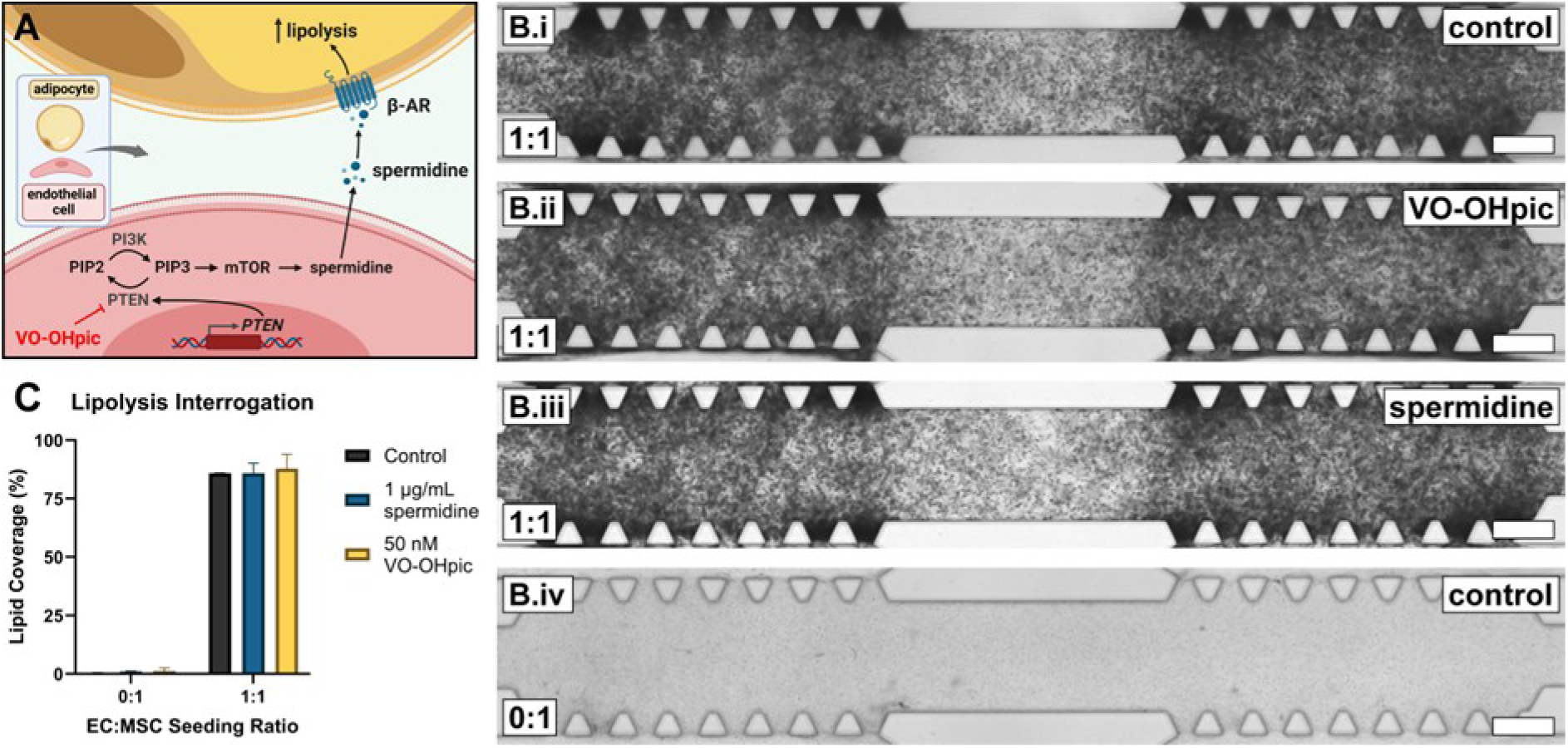
Proven inducers of lipolysis do not inhibit lipid formation in GFP-hAEC and hAD-MSC gradient co-cultures. **A.** A schematic representing the interactome of vascular PTEN-mediated lipolysis proposed by Monelli et al [4], the PTEN-inhibiting effect of VO-OHpic and lipolysis-enhancing effect of spermidine. Created with BioRender.com. **B.** Monochromatic brightfield images of day 31 1:1 seeding ratio cultures without inhibiting supplements (i) with 50 nM VO-OHpic (ii) and 1 µM spermidine (iii); and 0:1 seeding ratio cultures without inhibiting supplements (iv, scale bars = 500 µm). **C.** Quantitative analysis of lipid coverage across the culture compart after 31 days gradient culture comparing seeding ratios of 0:1 and 1:1 (N = 3 biological replicas for spermidine and for VO-OHpic, N = 2 control replicates, mean ± standard deviation). β-AR: beta-adrenaline receptor, PI3K: phosphoinositide 3-kinase. PIP2: phosphatidylinositol 4,5-bisphosphate, PIP3: phosphatidylinositol (3,4,5)-trisphosphate, mTOR: mammalian target of rapamycin, PTEN: phosphatase and tensin homolog, VO-OHpic: hydroxyl(oxo)vanadium 3-hydroxypiridine-2-carboxylic acid, EC: endothelial cell, MSC: mesenchymal stem cell.

## 4. Discussion

For the first time in scientific literature, we detail a thorough optimisation and validation of the culture conditions required to construct highly reproducible microvascular networks from green fluorescent protein-expressing hTERT immortalised human aortic endothelial cells (GFP-hAECs) co-cultured with hTERT immortalised adipose-derived mesenchymal stem cells (hAD-MSCs). Immortalised cell lines exhibit distinct advantages over their primary counterparts, which suffer from batch-to-batch variability and limited *in vitro* lifespan, both of which are detrimental to the reproducibility of microvascular network studies and their use for human preclinical models.

Endothelial (GFP-hAEC) to stromal cell (hAD-MSC) seeding ratio was a critical determinant of vascular network morphology, consistent with prior studies [64]. Seeding ratios of 2:1 and 1:1 resulted in entire culture monolayer detachment from the culture surface before day 7 (video S1). It is suspected that the formation of microvascular networks introduces tension on the cell monolayer, which is proportional to the number of endothelial cells. Additionally, these higher endothelial cell seeding ratios resulted in larger vessel junctions and reduced extravascular space (**figure 1.A.i-ii**), a morphology less consistent with *in vivo* human capillary networks [65]. The latter phenomenon was also evident, however, less so, in the 1:2 condition (**figure 1.A.iii**).

The lowest GFP-hAEC and hAD-MSC seeding ratio studied (1:10) resulted in the lowest vessel area and total vessel length (**figure 1.B.iii-iv**), almost certainly due to the reduced number of seeded endothelial cells. Interestingly, after day 4, the 1:10 GFP-hAEC:hAD-MSC condition demonstrated significant angiogenesis toward vacant extravascular space, evident both qualitatively in images and quantitatively by an uptick in all analysed vessel metrics (**figure 1.D**, figure S4). This knowledge is pertinent for future studies examining angiogenesis post-establishment of stable microvascular networks. The 1:5 condition demonstrated a maximum network density with non-enlarged junctions while maintaining surface adhesion throughout the 7-day culture period (**figure 1.A.iii and B, video S1**) and was therefore determined to be the optimal cell seeding ratio for 2D vasculogenesis. While both cell types were presumed to be homogeneously distributed during the seeding process, unevenness in surface attachment distribution was observed to result in a distribution in network density across the culture surface. This variability was accounted for by sampling fields of view across three technical replicates per each of three biological replicates.

Microvascular network phenotype was assessed against a panel of markers consisting of the endothelial cell marker, CD31; the pericyte-like cell marker, αSMA; the MSC marker, CD90; and the filamentous cytoskeletal protein marker, F-actin. As expected, the detection of CD31 was abundant in regions consistent with hAEC-GFP expression and vessel-like morphology (**figure 1.E.ii-iii**). At high magnification, localisation of CD31 detection appeared most abundant at cell-cell junctions in vessels, consistent with *in vivo* human vessel phenotype [66]. Detection for αSMA was observed across the culture surface, however, appeared most abundant in cells surrounding vessels (**figure 1.E.iv**). At high magnification, αSMA positive cells can be seen lining the outside of vessels, and, consistent with *in vivo* adipose-derived MSC or pericyte-like behaviour (figure S7.B.i) [67, 68] appear to be forming bridges across vessels (figure S7.B.iii), potentially as a precursor to angiogenesis and end-to-end anastomosis. The MSC marker CD90 was observed in cells across the culture, both on vessels as mural-like cells and between vessels on the 2D culture plastic surface, adopting an aligned morphology (**figure 1.E.v**). The retention of a CD90 phenotype indicates MSC multipotency, among other traits [69].

Time-lapse microscopy and image analysis was utilised to quantify vasculogenesis and characterise this morphogenetic process. Imaging typically commenced 2 hours after cell seeding, to ensure the movement of the microscope stage did not influence cell attachment, Additionally, a single technical replicate commencing imaging directly after seeding was conducted in isolation to confirm the morphogenetic behaviour during the 0–2 hours period (figure S3). All seeding ratios demonstrated a similar trend in total vessel area, total vessel length, number of junctions and number of end points, which was a sharp increase in magnitude from seeding to a peak at approximately 12 hours post-seeding (**figure 1.D**). This was then followed by a decrease in magnitude until approximately day 3 and followed by stabilisation until the termination of culture at day 7. It is therefore hypothesised from these observations that *in vitro* vascular morphogenesis occurs in three distinct morphogenetic phases: (1) the formation of discrete and discontinuous early vessel-like structures (0–0.5 days), (2) anastomosis of discrete early vessels towards a continuous network (0.5–4 days) and (3) maturation and readjustment of a continuous vessel network (4–7 days). In low-endothelial cell number cultures (e.g., 1:10 condition), angiogenesis can also be added to the third phase of development (figure S4). These findings also agree with the mechanisms reviewed by Venkatakrishnan *et al,* which describe the *de novo* assembly of discrete endothelial cells into the first blood vessels of the early embryo [70]. Both angiogenic and end-to-end anastomotic-like processes were observed via time lapse microscopy to occur in these cultures (figure S5), highlighting the utility of this system, not only for predictably studying vasculogenesis *in vitro,* but also angiogenesis and anastomosis. The removal of spent media and the addition of fresh media was observed to coincide with a small uptick in all measured vessel network properties (**figure 1.D, figure S3**), most likely due to the refreshing of active vasculogenic factors in the culture volume. The ability to accurately and quantitively assess and predict microvascular tissue development is of tremendous importance toward industrial standardisation of preclinical vascular models and therapeutic screening platforms. Furthermore, the ability to quantify *in vitro* tissue development in real time would allow informed and timely intervention to rescue, terminate, feed, differentiate or harvest a culture, improving process efficiency and reducing running costs for regenerative therapy manufacturing.

GFP-hAEC *in vitro* vasculogenesis was shown to critically rely on both the presence of hAD-MSCs in culture and the inclusion of an endothelial cell growth factor cocktail in media (figure S9). It was hypothesised that GFP-hAEC vasculogenesis could potentially proceed in the absence of an endothelial cell growth factor cocktail (and rely purely on paracrine signalling from hAD-MSCs) or in the absence of supporting hAD-MSCs, to aid simplicity and reduce culture cost. In the absence of the endothelial cell growth factor cocktail, GFP-hAECs did not attach to the culture substrate post-seeding (figure S9.B). Additionally, in the absence of supporting hAD-MSCs (in the presence of endothelial cell growth factor cocktail), GFP-hAECs simply attached and continued expanding as an overconfluent monolayer (figure S9.A). Interestingly, the metabolic activity of hAD-MSC cultures in complete endothelial cell media at days 5 and 7 of cultures was found to be more than double of that in complete MSC media (figure S8), potentially due to the presence of IGF-1, FGF, and/or heparin in complete vascular cell media, which are known promoters of MSC proliferation [71]. This potentially contributes to the activity of hAD-MSCs in vasculogenic co-culture, which was typically conducted in complete vascular cell media. Future analyses of growth factor usage, such as by ELISA or cytokine bead assays, would complement these media deprivation studies.

This vasculogenesis co-culture technology was translated to a fibrin hydrogel-supported 3D culture system within a commonly adopted microfluidic device configuration (figure S10) [28, 61, 62, 72]. The small molecule, aprotinin, was included in the hydrogel formulation to prevent the degradation of fibrin due to its antifibrinolytic properties [73]. Like 2D findings, GFP-hAEC and hAD-MSC co-cultures exhibited stable vessel network formation after 7-days culture in complete endothelial cell media and similarly exhibited a cell seeding ratio dependence on vascular network morphology (**figure 2.A-B**). At high seeding ratios (5:1, 2:1) vessel junctions increased in size and vessels merged into an agglomerated structure, subsequently causing an increase in vessel area and decreases in vessel length, number of junctions and number of end points (**figure 2.A.i-ii**). At the lowest seeding ratio (1:2), vessel networks were highest in length, junctions and end points, however, exhibited discontinuities throughout the network (**figure 2.A.iv**). A seeding ratio of 1:1 resulted in network continuity with maximum length, junctions and end points (**figure 2.A.iii**) and was determined to be the ideal seeding ratio for creating stable and robust vessel networks from these immortalised cell types suspended in fibrin hydrogels. Of note is the small magnitude of standard deviations of vascular metrics for both 3D and 2D vasculogenesis systems (**figure 1.B and 2.B**), indicating high reproducibility.

Consistent with 2D morphogenetic study outcomes, vessel networks formed in 3D culture environments (1:1 seeding ratio) demonstrated a rapid increase in junctions, vessel length and end points (0–0.5 days), followed by a decrease (0.5–3.5 days) and plateau (3.5–7 days). Unlike the 1:10 seeding ratio in 2D, no angiogenesis was observed during the plateau phase (**figure 2.D**). An identical immunocytochemical marker panel demonstrated similar detection of CD31 localised to vessel cell junctions, αSMA and CD90 in cells both on and between vessels and F-actin on the cytoskeleton of cells in vessels and in extravascular space (**figure 2.E**). Furthermore, confocal microscope image cross-sections confirmed vessel patency (**figure 2.F**). Unlike 2D cultures, αSMA was detected with similar intensity in cells occupying the extracellular space and cells lining vessels (**figure 1.E.iv** and **2.E.iv**). This difference is potentially a result of the increased dimensionality of the culture, or mechanical properties of the hydrogel influencing MSC differentiation towards pericyte-like phenotypes, evidenced by positive aSMA staining [74].

Similar to 2D cultures, 3D vasculogenesis was investigated in the absence of hAD-MSCs or in EC basal media (without a growth factor cocktail). Interestingly, unlike 2D results, vascular networks formed in both deprived 3D conditions (figure S14). In the absence of hAD-MSCs, endothelial cells still self-assembled into 3D vessel networks, as previously found [75], but network quality was poor, exhibited by discontinuities throughout the network and rough vessel borders (figure S14.A), likely a result of both absent paracrine signalling and the physical stabilising effects of pericyte-like cells, in the form of hAD-MSCs. In the absence of endothelial cell media growth factor cocktail, GFP-hAEC and hAD-MSC co-cultures still formed continuous 3D vessel networks with tight vessel borders, however, with smaller vessel diameter and junctions compared to those with the growth factor cocktail (figure S14.B). These differences suggest that GFP-hAECs and hAD-MSCs provide mutually supportive paracrine factors, such as MSC-derived VEGF [76], enabling microvascular network establishment in 3D (but less so 2D). Additionally, these findings suggest vessel and junction diameters (and therefore projected area) could be tuned based on the concentration of growth factors in basal media.

The presence of vessel openings spanning inter-pillar spacings is a phenomenon sometimes seen in vasculogenic co-cultures using similar microfluid device configurations with alternative cell types, such as human umbilical vein endothelial cells [61, 72]. Vessel networks in this study did not form visible or functional openings between pillars (figure S15.C), which closure was confirmed via addition of fluorescently labelled dextran (data not shown). To encourage vessel openings between pillars, an approach similar to that of Wan *et al* [28] was adopted, whereby, 13 × 10^6^ GFP-hAECs/mL (no hAD-MSCs) were seeded into the device and immediately aspirated, leaving a suspension of entirely endothelial cells between the pillars (figure S15.A), after which, the 1:1 co-culture was used to fill the culture compartment completely. While this approach aided in providing broader vessel structures between pillars (figure S15.B), these structures remained closed to the neighbouring media channel and did not allow the passage of fluorescently labelled dextran (data not shown). The reason as to why this model did not support typically vessel opening between pillar spacings is unknown. One hypothesis is that mural cell-like contractility prevents vessel opening, as the downregulation of CD90 in hAD-MSCs was demonstrated in late passage hTERT immortalised human fibroblasts by Wan *et al* to inhibit the ability of forming inter-pillar openings [77], however, this was not actively examined in this study.

To promote 3D co-culture vasculogenesis simultaneously with adipogenesis, a microfluidic chip was designed to create hydrogel gradients of stromal expansion, vasculogenic, and/or adipogenic growth factors (**figure 3.B**). This design was informed by growth factor diffusion-decay using partial differential equations across a simplified 2D cartesian model of microchip hydrogel space [31]. Every 2 days, growth factors are supplied through media channels on four edges of the microchip. The factors continuously decay in the media channel, and as they diffuse into the hydrogel, activating cells when above an effective concentration (EC50). While a useful design tool, our simulations assume growth factors diffuse and decay similarly as found in liquid cell culture media, and do not consider how cellularised fibrin hydrogels release, bind, or otherwise alter growth factor kinetics [34]. These growth factor assumptions should be experimentally validated or corrected through tracking fluorescently conjugated growth factors or sugars or measuring growth factor concentration found within opposite media channels. Nonetheless, this model predicts that the microfluidic chip design (**figure 3.B**) forms growth factor gradients above and below bioactive EC50 concentrations across the length of the culture compartment, for vasculogenic, stromal, and adipogenic factors (**figure 3.C, figure S17**).

When differentiated using a gradient culture device (**figure 3**), vessel networks and adipocytes were observed to form within the same 3D space (**figure 4.A**). The adipogen-rich side of the device was not found to inhibit the formation of continuous vascular networks, which were observed throughout the culture compartment after day 17. Post-inversion of culture gradients, at day 31, vessel network morphology was maintained and lipid droplet intensity increased (**figure 4.A, figure S18**), however, full homogeneity of lipid production across the culture compartment was not entirely achieved (**figure 4.A.ii, F.ii and G.ii**). It is hypothesised that hAD-MSCs lose their differentiation potential when maintained in a vasculogenic environment for 17 days [78]. The intensity of lipid production in the presence of microvascular networks was remarkably high (67.4 ± 0.07%) when compared to hAD-MSC 3D monocultures (1.86 ± 1.09%), consistent with the landmark murine model findings of Rupnick *et al* [79]. The intensity of lipogenesis was surprising given the limited adipogenic potential of hTERT immortalised hAD-MSCs reported by Masnikov *et al* [80]. While the mechanisms which underpin this vasculo-adipo relationship have remained poorly understood, Monelli *et al* reported a mechanism, proposing vessel endothelium aids in suppressing adipose tissue lipolysis [4, 81, 82]. By deleting endothelial *Pten* (phosphatase and tensin homolog) in mice, Monelli and colleagues observed a reduction in bodyweight and adiposity compared to control littermates. They confirmed *Pten* deletion supported sustained phosphoinositide 3-kinase (PI3K) activity and enhanced polyamine synthesis (specifically spermidine) in endothelial cells, which promoted beta-adrenaline receptor activity and upregulated lipolysis in *ex vivo* mouse WAT explants, primary mouse adipocytes and *in vivo* murine models. This led us to investigate whether this PTEN regulated mechanism could regulate lipid production in our human 3D co-culture model.

VO-OHpic is a known inhibitor of PTEN, with reported EC50 value of 50 nM [83, 84] and as described above, spermidine, a polyamine, has been demonstrated to induce lipolysis in primary mouse adipocytes at a concentration of 1 µM [4, 85, 86]. These two molecules at these concentrations were therefore chosen to investigate their ability of inducing lipolysis in our human co-culture model. Surprisingly, supplementation of this co-culture differentiation model with said compounds resulted in no change in percentage lipid coverage after 31-days co-culture (**figure 5, figure S18**). It is hypothesised that the limited complexity or maturity of this simple co-culture model does not completely recapitulate the multifaceted interactions of multicellular murine WAT tissue explants or the maturity of primary adipocytes. Or, potentially, the selected concentrations of the chosen molecules were insufficient to illicit the expected biological responses in immortalised human cell lines. Our human culture model deviates in some aspects from peripheral adipose tissue anatomy which contains fewer vessels of smaller diameter, more like figure S14.B [87, 88]. In addition, recent clinical trials have suggested that human spermidine pharmacokinetics vary from those proposed in murine models [89, 90].

Inducing the formation of physiologically relevant microvascular networks within developing *in vitro* tissue constructs is challenging due to the inherent nature of nutrient delivery in modern 2D batch culture approaches, which fail to mimic the complex spatial and temporal delivery seen *in vivo*. Technology which supports reproducible spatial and temporal control of biochemical growth environments is necessary for the development and subsequent scale up of *in vitro* tissue and organ products. By designing 2D, 3D hydrogel, and microfluidic co-culture technology to control spatiotemporal processes of vasculogenesis, we deliver a highly reproducible diagnostic system using immortalised human cell lines. We then apply our 3D hydrogel microfluidic model to generate counter-current media gradients, demonstrating a new approach for the co-development and integration of MSC-derived adipocytes and patent microvascular networks, within the same 3D tissue space. This serves as a platform technology for the scale up of vascularised mature connective tissue co-cultures, potentially adopting 2D and 3D channel arrays to control growth and differentiation in 3D, providing future avenues in vascular tissue models and for disease and biotherapeutic screening.

## 5. Conclusion

Biomanufacturing of vascularised connective tissue has broad applications towards tissue implant engineering, drug discovery, and cultivated meat. However, vasculogenic co-cultures suffer from reproducibility, scale-up to 3D, and an inability to vascularise and differentiate connective tissue simultaneously. In this study, optimal, robust and phenotypic vascular networks were formed in 2D culture by co-seeding hTERT immortalised green fluorescent protein-expressing human aortic endothelial cells (GFP-hAECs) with hTERT immortalised human adipose-derived mesenchymal stem cells (hAD-MSCs) at a density of 187,500 total cells/cm^2^ and a seeding ratio of 1:5. GFP-hAECs and hAD-MSCs similarly formed robust and phenotypical vascular networks when seeded suspended in 3D fibrin hydrogels at a density of 13 × 10^6^ total cells/mL with a seeding ratio of 1:1. Time-lapse microscopy and quantitative image analysis revealed a consistent trend in vessel network junctions, end points and length throughout development, demonstrated by an initial sharp rise, peak, decrease and plateau in morphological properties. Gradient microfluidic culture supported the co-formation of integrated microvascular networks and functional adipocytes within a single 3D hydrogel structure. Adipogenesis was drastically enhanced in the presence of vessel networks (67.4% lipid coverage) compared to in their absence (1.86% lipid coverage). Supplementation of gradient co-cultures with 50 nM VO-OHpic and 1 µM spermidine did not suppress lipogenesis, as reported with murine models. This research demonstrates a new *in vitro* culture system capable of generating reproducible vascularised adipose tissue from existing commercial immortalised human EC-and MSC-lines. This model has significant potential for studying vascular-adipose crosstalk in human preclinical studies of diabetes, obesity, and for biotherapeutic screening.

## Acknowledgements

This work was supported by an Advance Queensland Fellowship to MCA (AQIRF1312018), an Australian Research Council DECRA Fellowship to MCA (DE220100757), and a Ramaciotti Health Investment Grant from Perpetual Philanthropies to MCA (2022HIG08). This work used the Queensland node of the NCRIS-enabled Australian National Fabrication Facility (ANFF).

The authors would like to acknowledge insightful and productive discussions held with Associate Professor Joy Wolfram and Professor Justin Cooper-White.

## Credit Author Statement

The following authors contributed to the following CRediT (Contribute Roles Taxonomy) roles:

ARM: conceptualisation, data curation, formal analysis, investigation, methodology, resources, supervision, visualisation, writing - original draft, writing - review and editing.

RAF: formal analysis, investigation, writing - review and editing.

MCA: conceptualisation, formal analysis, funding acquisition, project administration, resources, software, supervision, writing - review and editing.

## Supplementary Information

**Table S1.** Primary and secondary antibody details.

| Supplier | Catalogue Number | Antigen | Species Reactivity | Host, Isotype | Stock Concentration (mg/mL) | Working Concentration (µg/mL) |
| --- | --- | --- | --- | --- | --- | --- |
| ThermoFisher Scientific | MA5-32559 | CD90 | Human | Rabbit, IgG | 1 | 10 |
| ThermoFisher Scientific | MA3100 | CD31 | Human | Mouse, IgG2a | 1 | 10 |
| ThermoFisher Scientific | MA5-15806 | aSMA | Human | Mouse, IgG1 | Not specified | (1:250 dilution) |
| ThermoFisher Scientific | MA5-14889 | ppARg | Human | Rabbit, IgG | 0.125 | 1.25 |
| ThermoFisher Scientific | 14-5698-82 | Ki67 | Human | Rat, IgG2a | 0.5 | 5 |
| ThermoFisher Scientific | 31235 | IgG isotype control | - | Rabbit, IgG | 11 | 10, 1.25 |
| ThermoFisher Scientific | 14-4724-82 | IgG2a isotype control | - | Mouse, IgG2a, kappa | 0.5 | 10 |
| ThermoFisher Scientific | 14-4714-82 | IgG1 isotype control | - | Mouse IgG1, kappa | 0.5 | (1:250 dilution) |
| ThermoFisher Scientific | A-11008 | Goat anti-Rabbit IgG (H+L), Alexa Fluor™ 488 | Rabbit | Goat, IgG | 2 | 4 |
| ThermoFisher Scientific | A-21434 | Goat anti-Rat IgG (H+L), Alexa Fluor™ 555 | Rat | Goat, IgG | 2 | 4 |
| ThermoFisher Scientific | A-21137 | Goat anti-Mouse IgG2a, Alexa Fluor™ 555 | Mouse | Goat, IgG | 2 | 4 |
| ThermoFisher Scientific | A-21245 | Goat anti-Rabbit IgG (H+L), Alexa Fluor™ 647 | Rabbit | Goat, IgG | 2 | 4 |
| ThermoFisher Scientific | A-21241 | Goat anti-Mouse IgG2a, Alexa Fluor™ 647 | Mouse | Goat, IgG | 2 | 4 |
| ThermoFisher Scientific | A-21240 | Goat anti-Mouse IgG1, Alexa Fluor™ 647 | Mouse | Goat, IgG | 2 | 4 |

---

**Figure S1.**
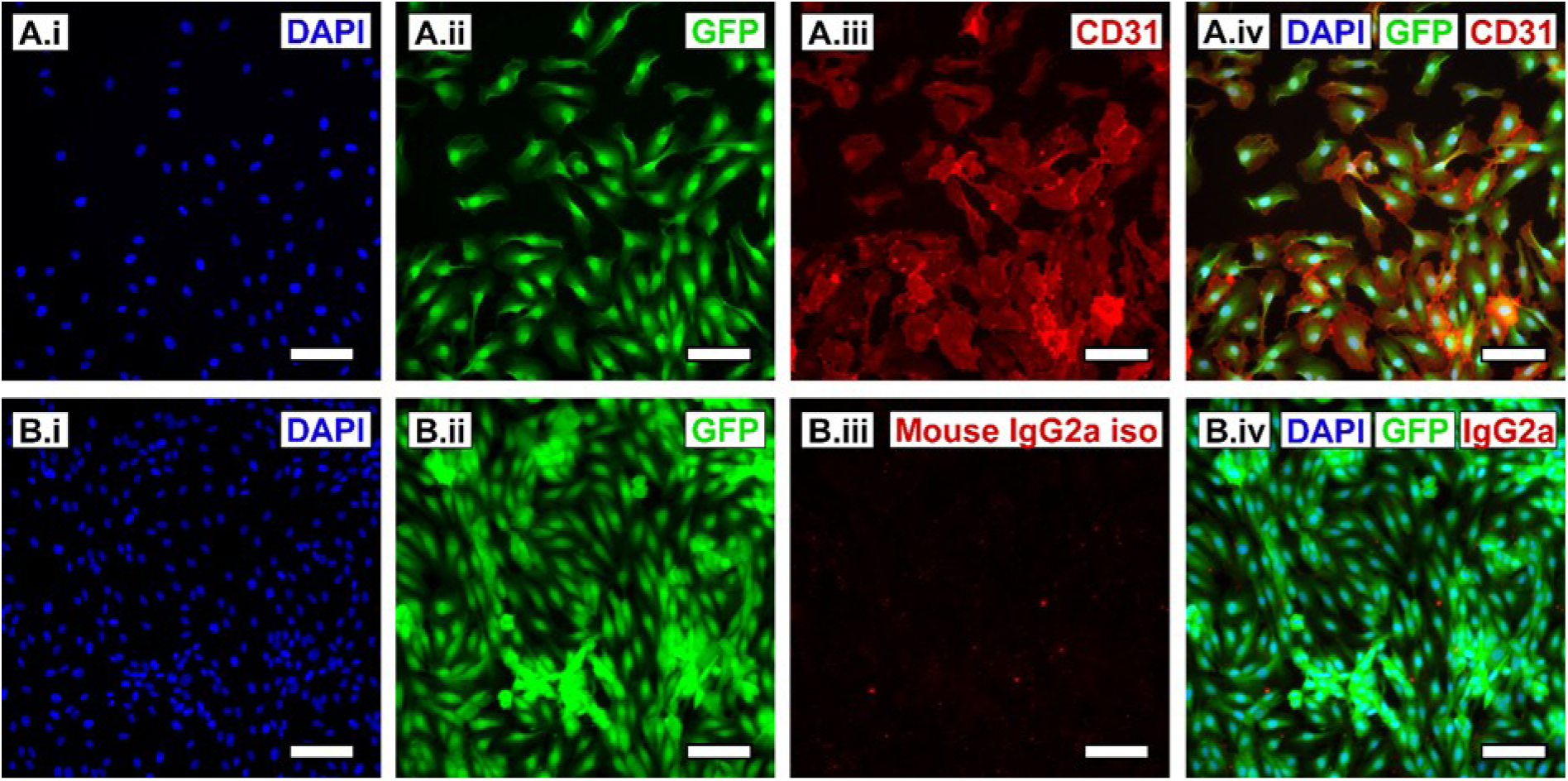
GFP-hAEC immunocytochemical (ICC) fluorescence characterisation. **A.** ICC detection for cluster of differentiation 31 (CD31) with images representing nuclear counterstain DAPI (**i**), endogenous GFP expression (**ii**), CD31 (**iii**) and a merge of the three (**iv**). **B.** ICC detection for the mouse IgG2a isotype control antibody with images representing nuclear counterstain DAPI (**i**), endogenous GFP expression (**ii**), mouse IgG2a isotype control (**iii**) and a merge of the three (**iv**). Scale bars = 100 µm. DAPI: 4’,6-diamidino-2-phenylindole, GFP: green fluorescent protein, CD31: cluster of differentiation 31, iso: isotype, M: mouse, IgG: immunoglobulin G.

**Figure S2.**
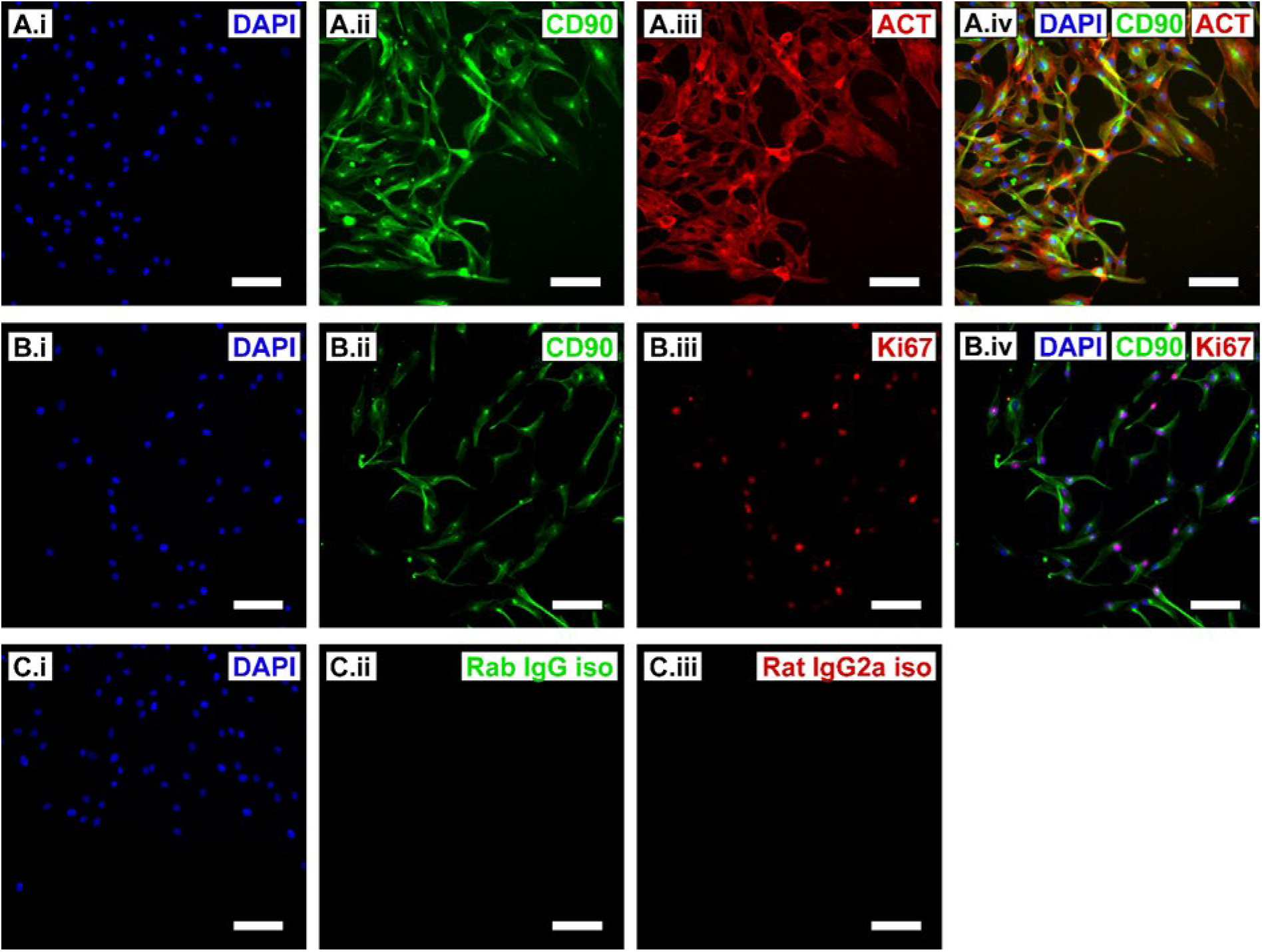
hAD-MSC immunocytochemical (ICC) fluorescence characterisation. **A.** ICC detection for cluster of differentiation 90 (CD90) with images representing nuclear counterstain DAPI (**i**, blue), CD90 (**ii**, green), filamentous actin (**iii**, red) and a merge of the three (**iv**). **B.** ICC detection for Kiel 76 (Ki67) with images representing nuclear counterstain DAPI (**i**, blue), CD90 (**ii**, green), Ki67 (**iii**, red) and a merge of the three (**iv**). **C.** ICC detection for isotype control antibodies with images representing nuclear counterstain DAPI (**i**, blue), rabbit IgG isotype (**ii**, green) and rat IgG2a isotype (**iii**, red). Scale bars = 100 µm. DAPI: 4’,6-diamidino-2-phenylindole, CD90: cluster of differentiation 90, IgG: immunoglobulin G, iso: isotype, rab: rabbit.

Videos are attached to the end of this bioRxiv submission.

Video S1. Time-lapse microscopy of GFP-hAECs and hAD-MSCs captures spontaneous formation of 2D microvascular networks with varying seeding ratio. Fluorescence time-lapse imaging (20 minute intervals) of GFP-hAEC microvascular networks formed over 7-days culture in complete vascular cell media, with GFP-hAEC:hAD-MSC seeding ratios of 2:1, 1:1, 1:2, 1:5 and 1:10 (scale bars = 1 mm).

**Figure S3.**
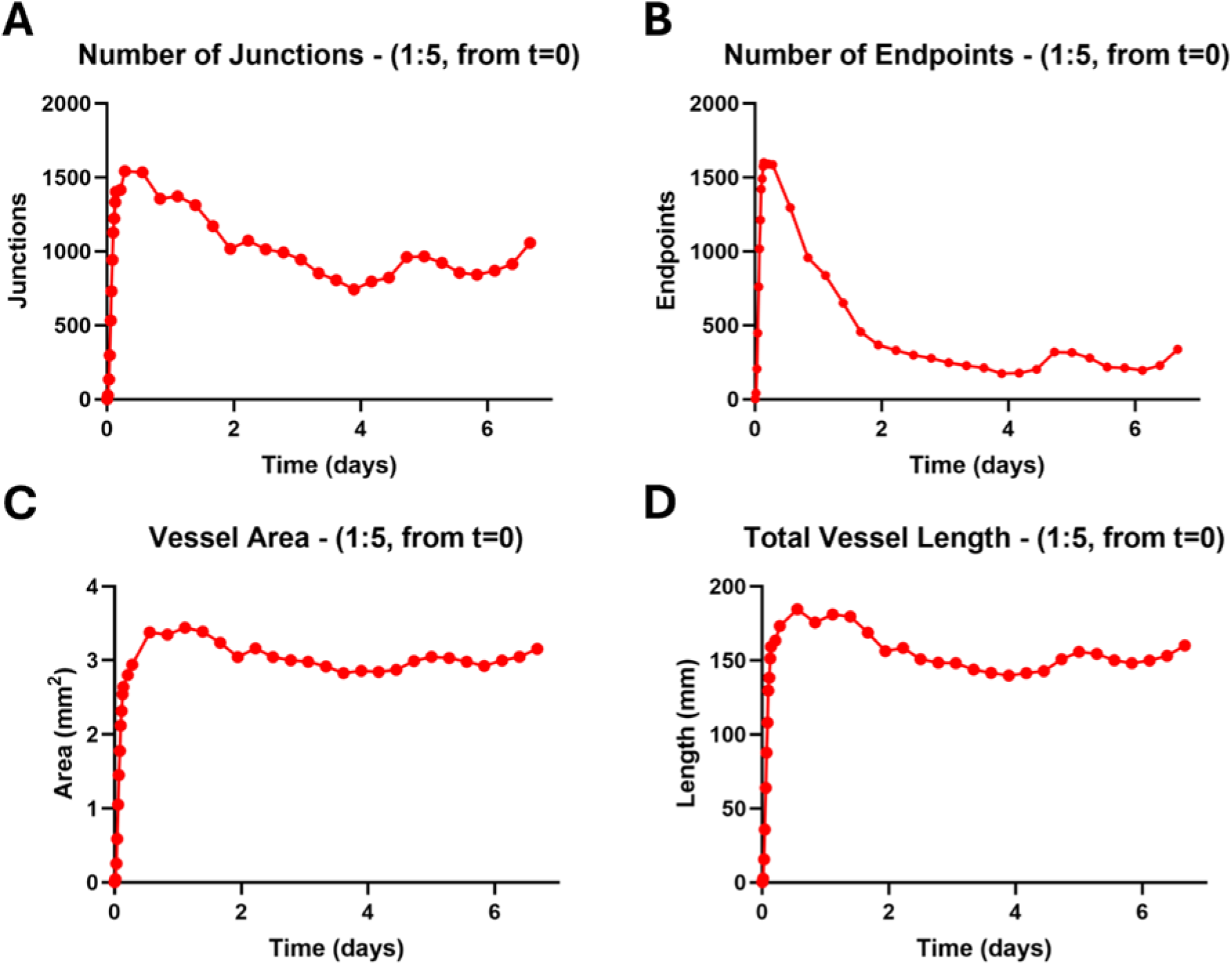
GFP-hAEC and hAD-MSC vasculogenesis co-culture time-lapse quantification. Quantitative morphological analysis of image set commenced directly post-seeding for the entire length of the culture. **A.** Number of vessel network junctions. **B.** Number of vessel endpoints. **C.** Total vessel network area. **D.** Total vessel network length.

**Figure S4.**
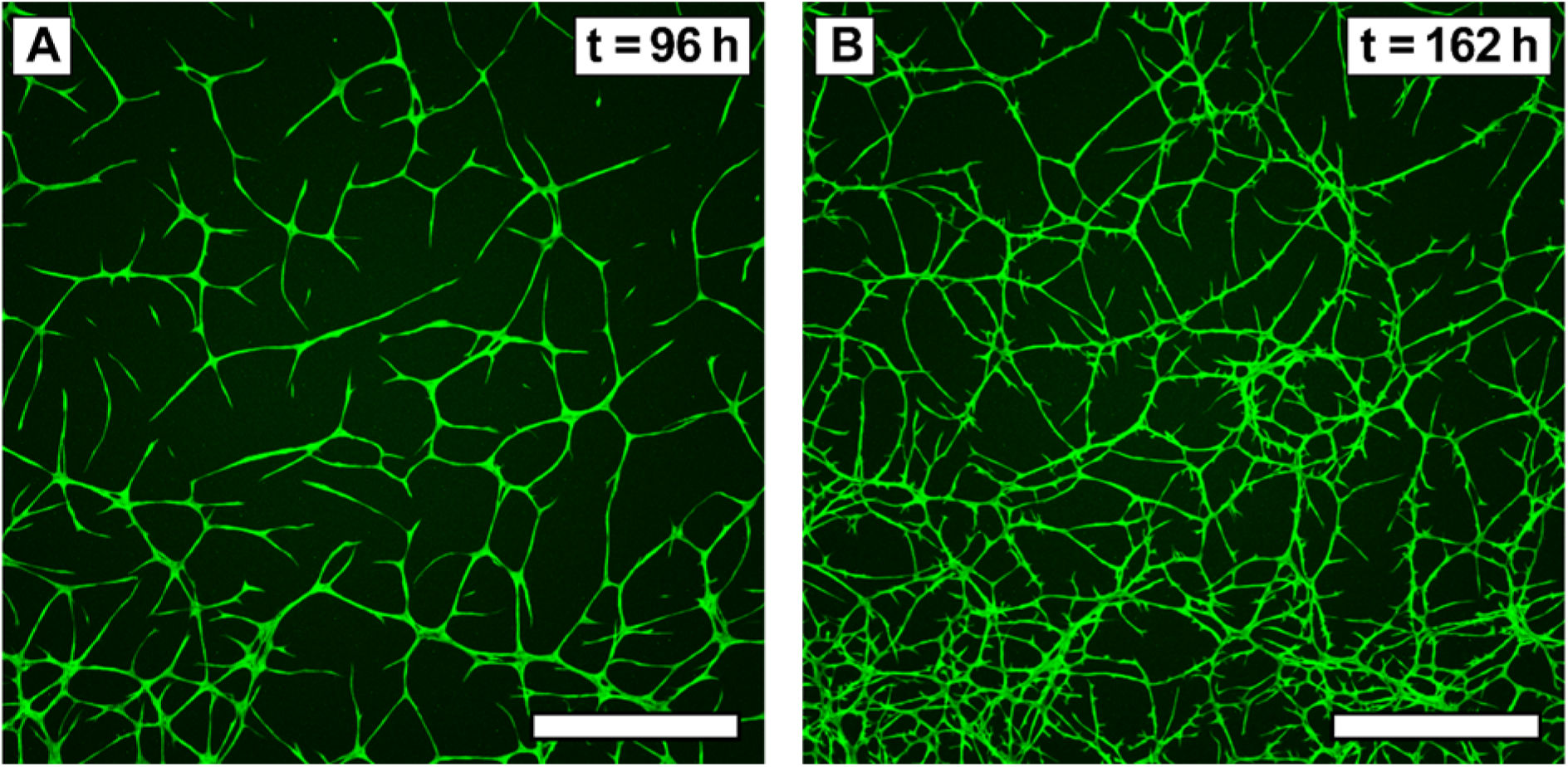
GFP-hAEC and hAD-MSC vasculogenesis 1:10 co-cultures exhibit angiogenesis-like behaviour in the latter stages of a 7-day culture. A. At the 96^th^ hour of culture, vessels display elongated morphology with no spine (scale bar = 1 mm). B. At the 162^nd^ hour of culture, vessels develop short spines extending from long continuous vessels (scale bar = 1 mm).

**Figure S5.**
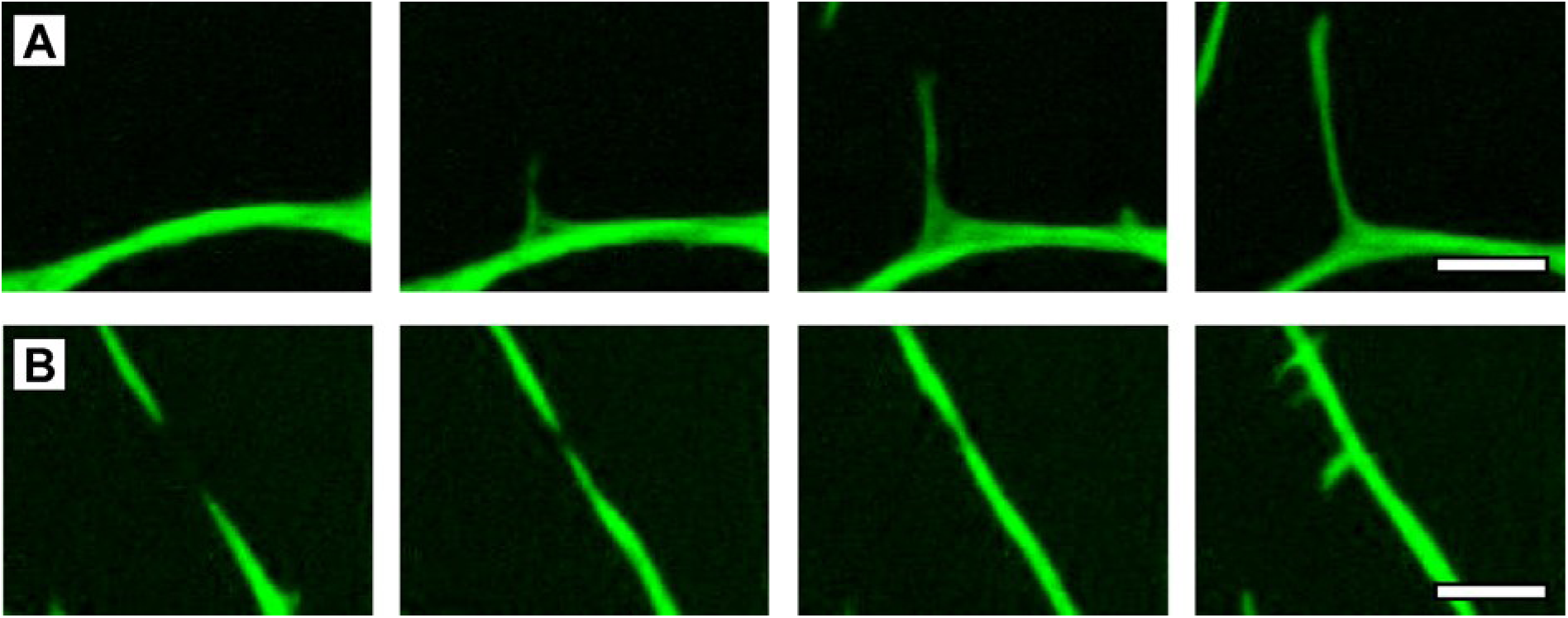
Vasculogenesis cultures replicate vessel morphogenetic processes other than vasculogenesis. **A.** Angiogenesis can be observed as the protrusion of a single tip from an existing vessel which grows in length over time (scale bar = 100 µm). **B.** End-to-end anastomosis can also be observed when two vessel tips grow towards each other and eventually fuse into a single continuous vessel (scale bar = 100 µm).

**Figure S6.**
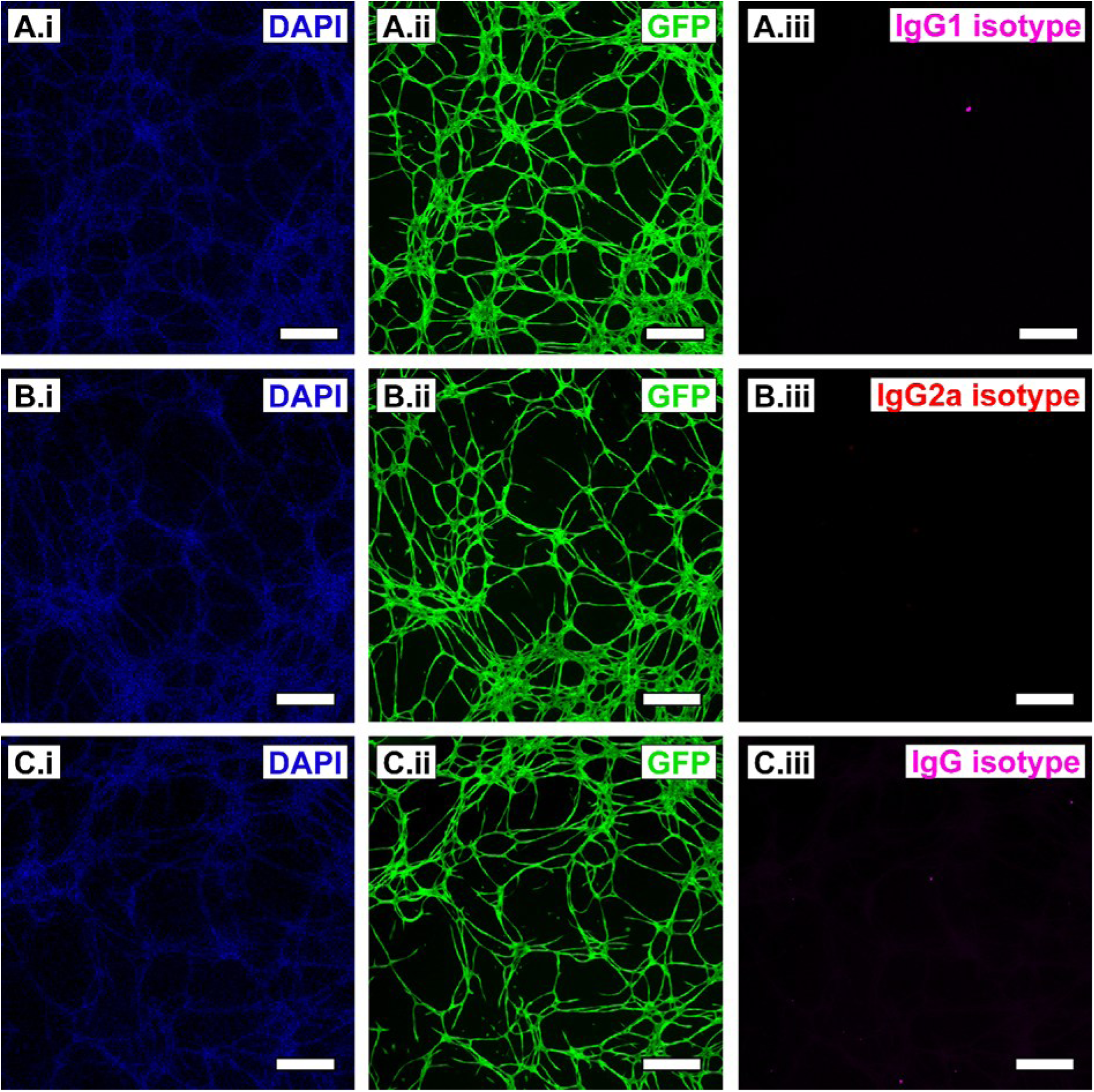
GFP-hAEC and hAD-MSC vasculogenesis co-culture day 7 (seeded at 1:5 ratio) immunocytochemical (ICC) fluorescence characterisation, isotype control stains. A. Mouse IgG1 isotype control with images representing nuclear counterstain DAPI (i, blue), GFP (ii, green), mouse IgG isotype control antibody (iii, magenta). B. Mouse IgG2a isotype control with images representing nuclear counterstain DAPI (i, blue), GFP (ii, green), mouse IgG2a isotype control antibody (iii, red). C. Rabbit IgG isotype control with images representing nuclear counterstain DAPI (i, blue), GFP (ii, green), rabbit IgG isotype control antibody (iii, magenta). Scale bars = 500 µm. DAPI: 4’,6-diamidino-2-phenylindole, GFP: green fluorescent protein.

**Figure S7.**
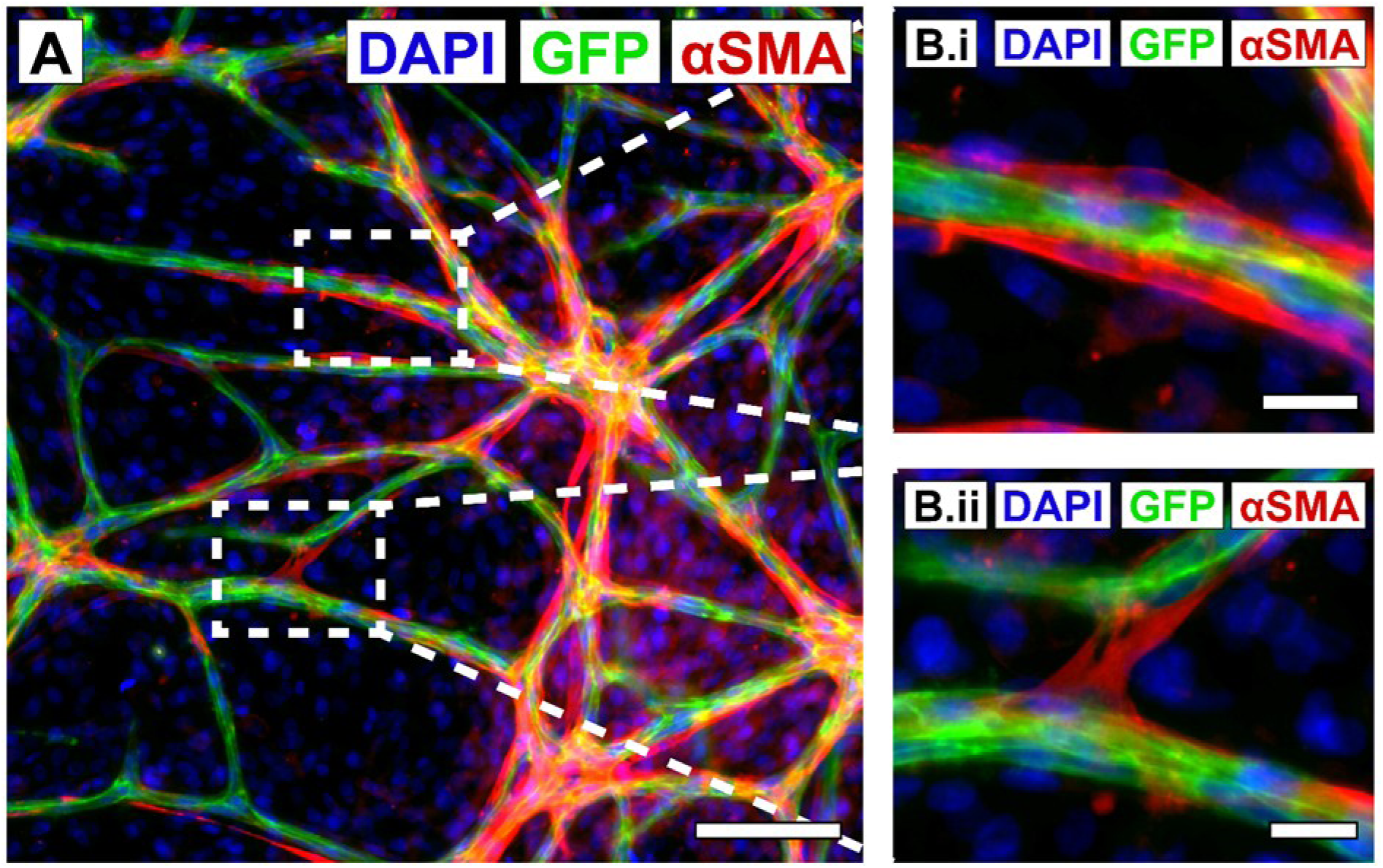
GFP-hAEC and hAD-MSC vasculogenesis co-cultures display pericyte-like behaviour. **A.** Vasculogenesis 1:5 cultures fixed at day 7 and immunocytochemically stained for alpha smooth muscle actin (αSMA), nuclear counter stained with DAPI and captured for endogenous GFP (scale bar = 100 µm). **B.** Cells positive for αSMA are observed lining the outside of vessels (i, scale bar = 20 µm) and forming bridges between vessels (ii, scale bar = 20 µm).

**Figure S8.**
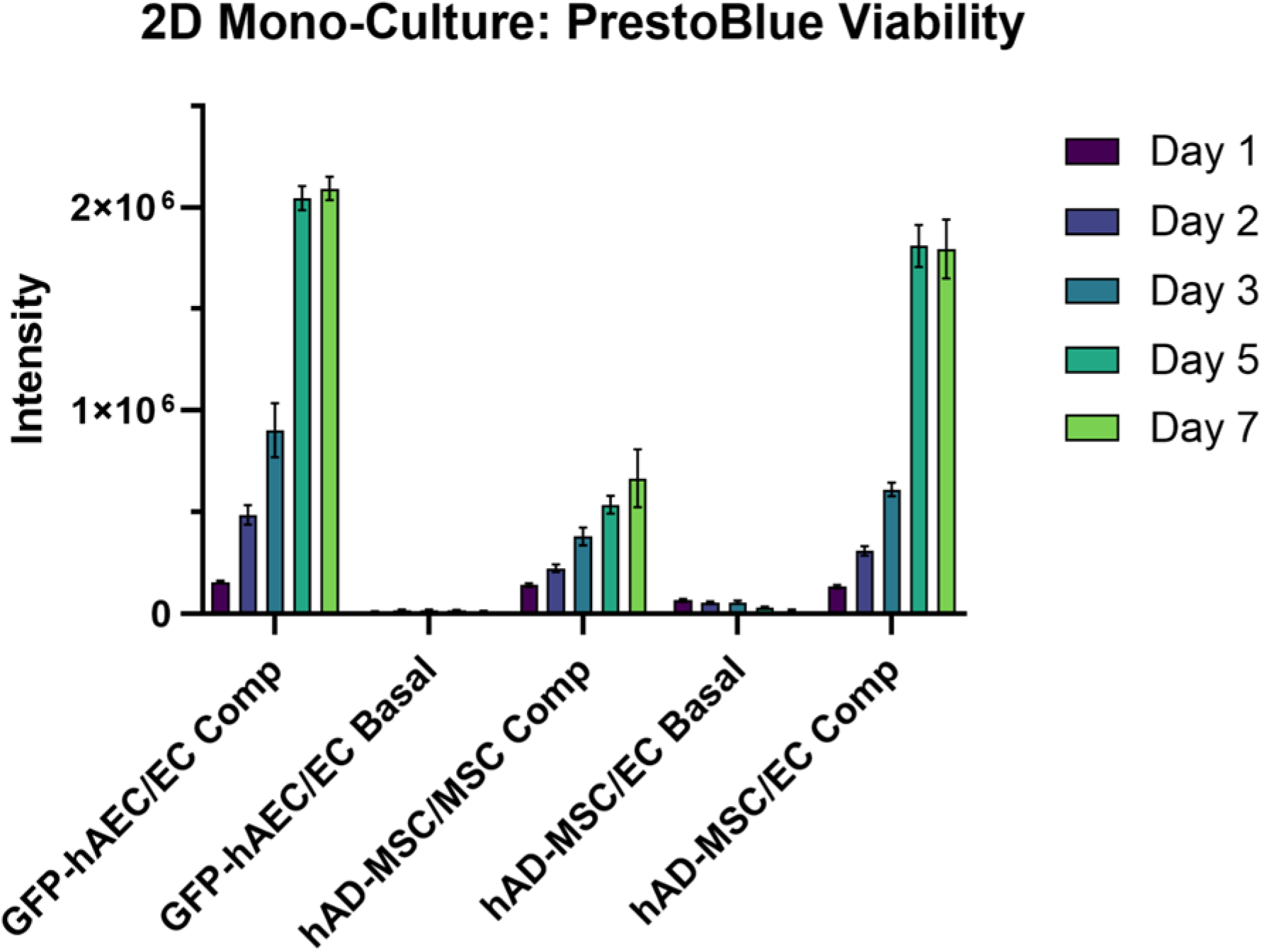
GFP-hAECs and hAD-MSCs demonstrate altered culture viability in the presence of different media compositions. GFP-hAECs were culture in complete vascular cell media and vascular cell basal cell media; and hAD-MSCs in complete MSC media, MSC basal media and complete vascular cell media. EC comp: complete vascular cell media, EC basal: vascular cell basal media, MSC comp: complete MSC media. (N = 3 technical replicas)

**Figure S9.**
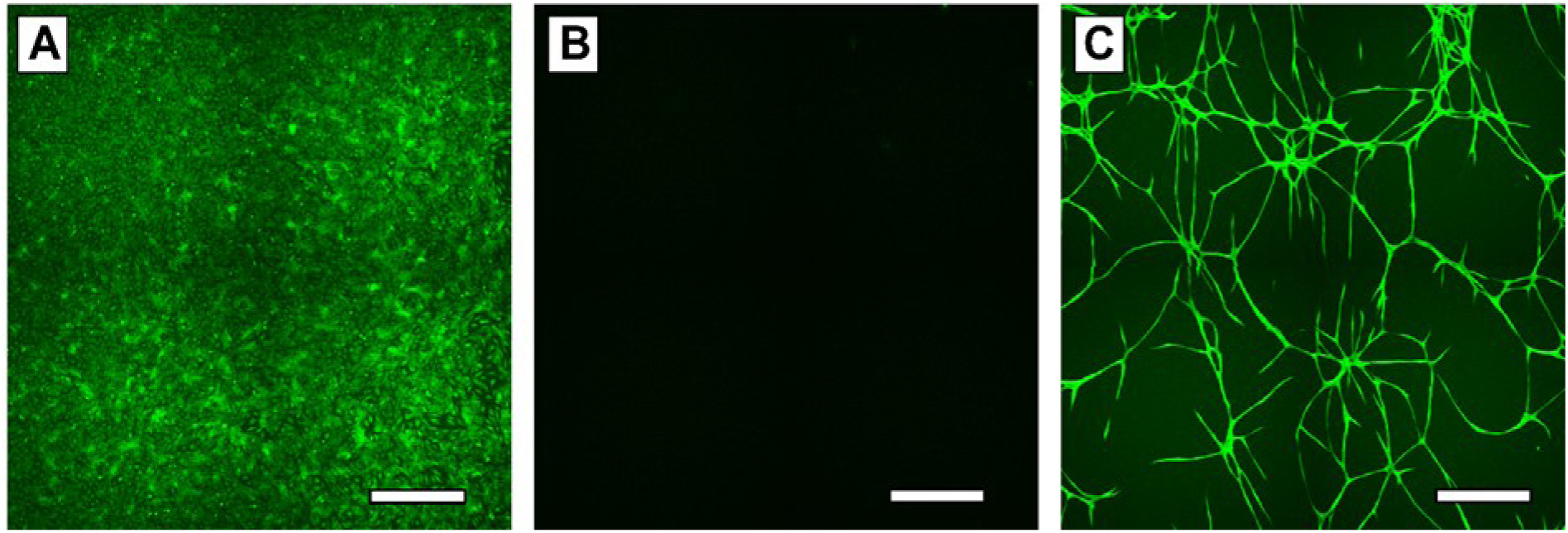
GFP-hAECs only form vessel networks in combination with endothelial cell growth factors and supporting hAD-MSCs. **A.** In the absence of hAS-MSCs, GFP-hAECs form a dense monolayer after 7-days culture (scale bar = 500 µm). **B.** In the absence of vascular cell growth factors, GFP-hAECs do not attach to the culture surface (scale bar = 500 µm). C. In the presence of hAD-MSCs and complete vascular cell media, GFP-hAECs spontaneously form vessel networks after 7-days culture (scale bars = 500 µm).

**Figure S10.**
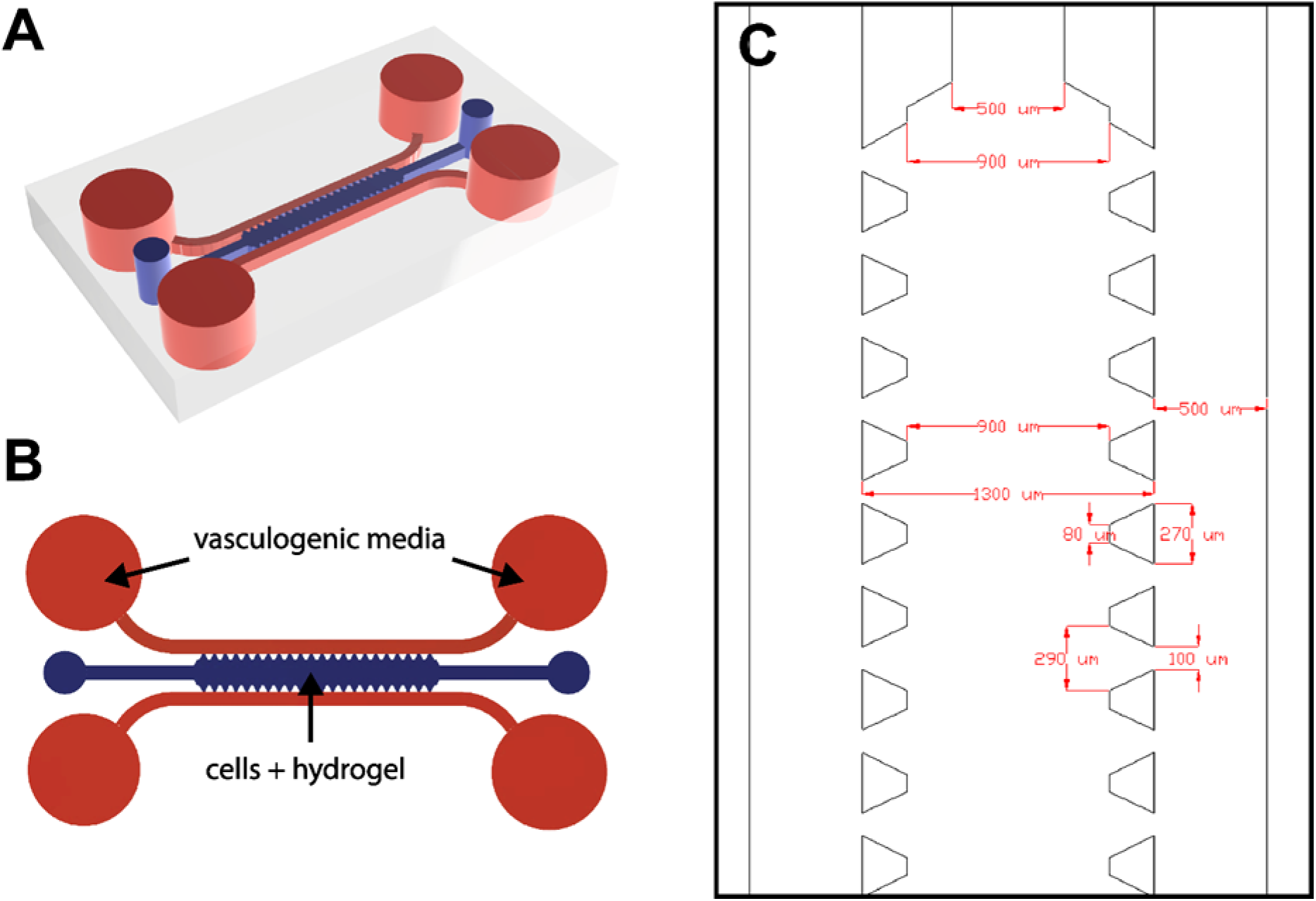
Microfluidic culture device for supporting 3D hydrogel embedded vasculogenesis. **A.** Device rendering. **B.** Device schematic depicting the cell-hydrogel culture compartment (blue) and flanking vasculogenic media channels (red). **C.** Device dimensions.

Videos are attached to the end of this bioRxiv submission.

**Video S2. Time-lapse microscopy GFP-hAECs and hAD-MSCs captures spontaneous formation of 3D microvascular networks.** Fluorescence (left) and brightfield (right) time-lapse imaging (30 minute intervals) of GFP-hAEC microvascular networks formed over 7-days culture in complete vascular cell media, with GFP-hAEC:hAD-MSC seeding ratios of 1:1 (field of view = 1.33 × 1.33 mm).

Videos are attached to the end of this bioRxiv submission.

**Video S3. Time-lapse microscopy and segmented image analysis outputs of GFP-hAECs and hAD-MSCs captures and quantified spontaneous formation of 3D microvascular networks.** Fluorescence (left) and overlayed segmentation and skeletonization (right) of time-lapse imaging (300 minute intervals) of GFP-hAEC microvascular networks formed over 7-days culture in complete vascular cell media, with GFP-hAEC:hAD-MSC seeding ratios of 1:1 (field of view = 1.33 × 1.33 mm).

**Figure S11.**
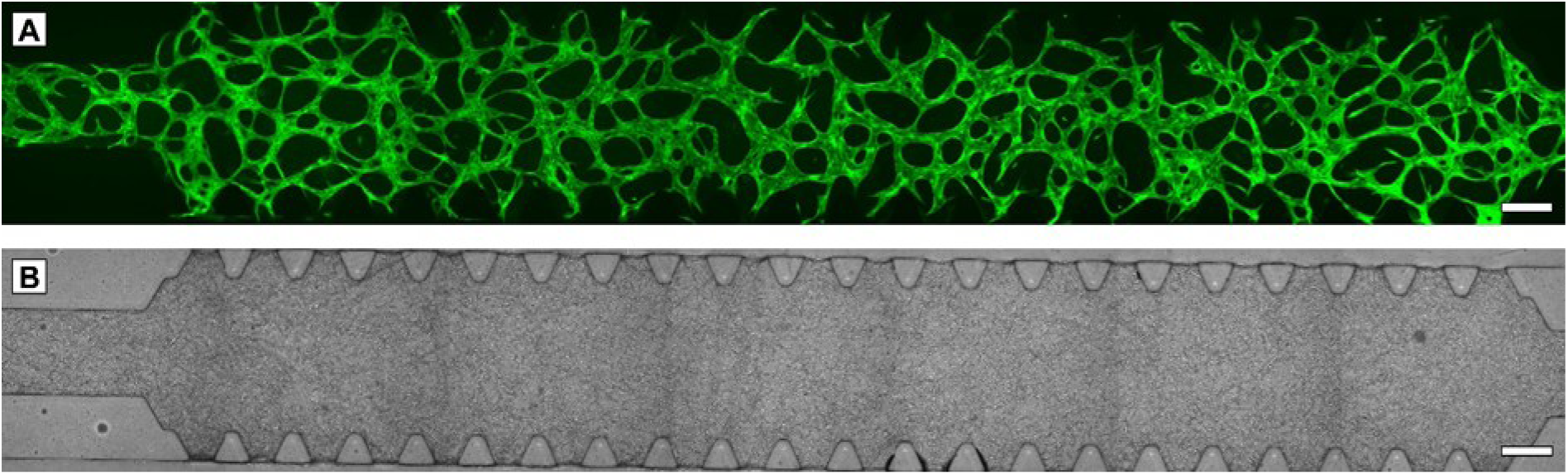
GFP-hAECs and hAD-MSCs spontaneously form robust 3D hydrogel-embedded microvascular networks across the entire length of microfluidic device culture compartments. Fluorescence (A) and Brightfield (B) images of GFP-hAECs as microvascular networks formed after 7-days fibrin hydrogel-embedded microfluidic device culture in complete vascular cell media with GFP-hAEC:hAD-MSC seeding ratios of 1:1 (scale bars = 300 µm).

**Figure S12.**
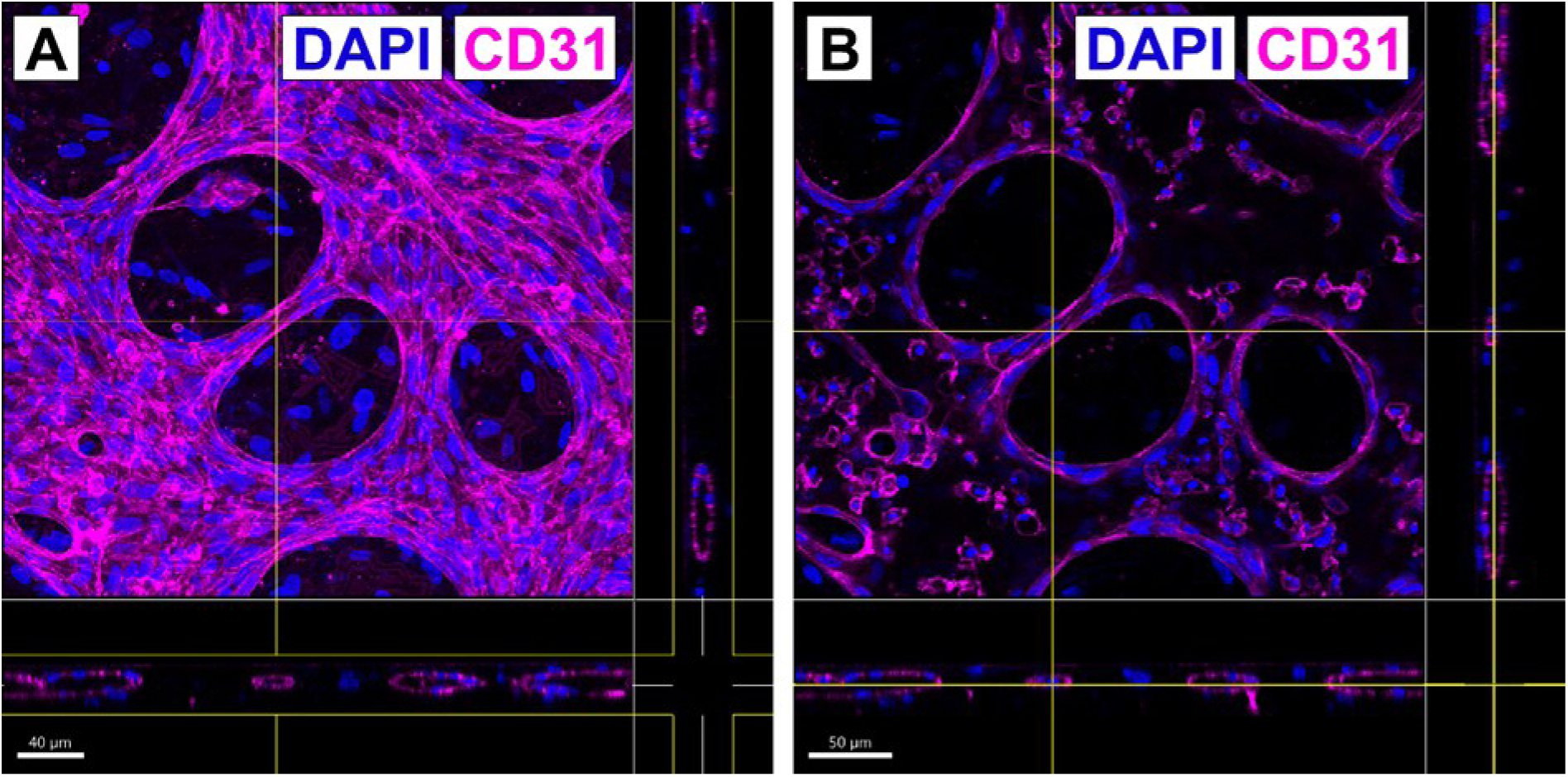
Single CD31 positive cells reside within microvascular network lumen. **A.** Maximum intensity projection of a CD31^+^ vessel network demonstrated vessel patency in the xz and yz cross-sectional plane (scale bar = 40 µm). B. Single round CD31^+^ cells can be seen suspended in the xy plane bisecting the vessel network (scale bar = 50 µm). DAPI: 4’,6-diamidino-2-phenylindole, CD31: cluster of differentiation 31.

**Figure S13.**
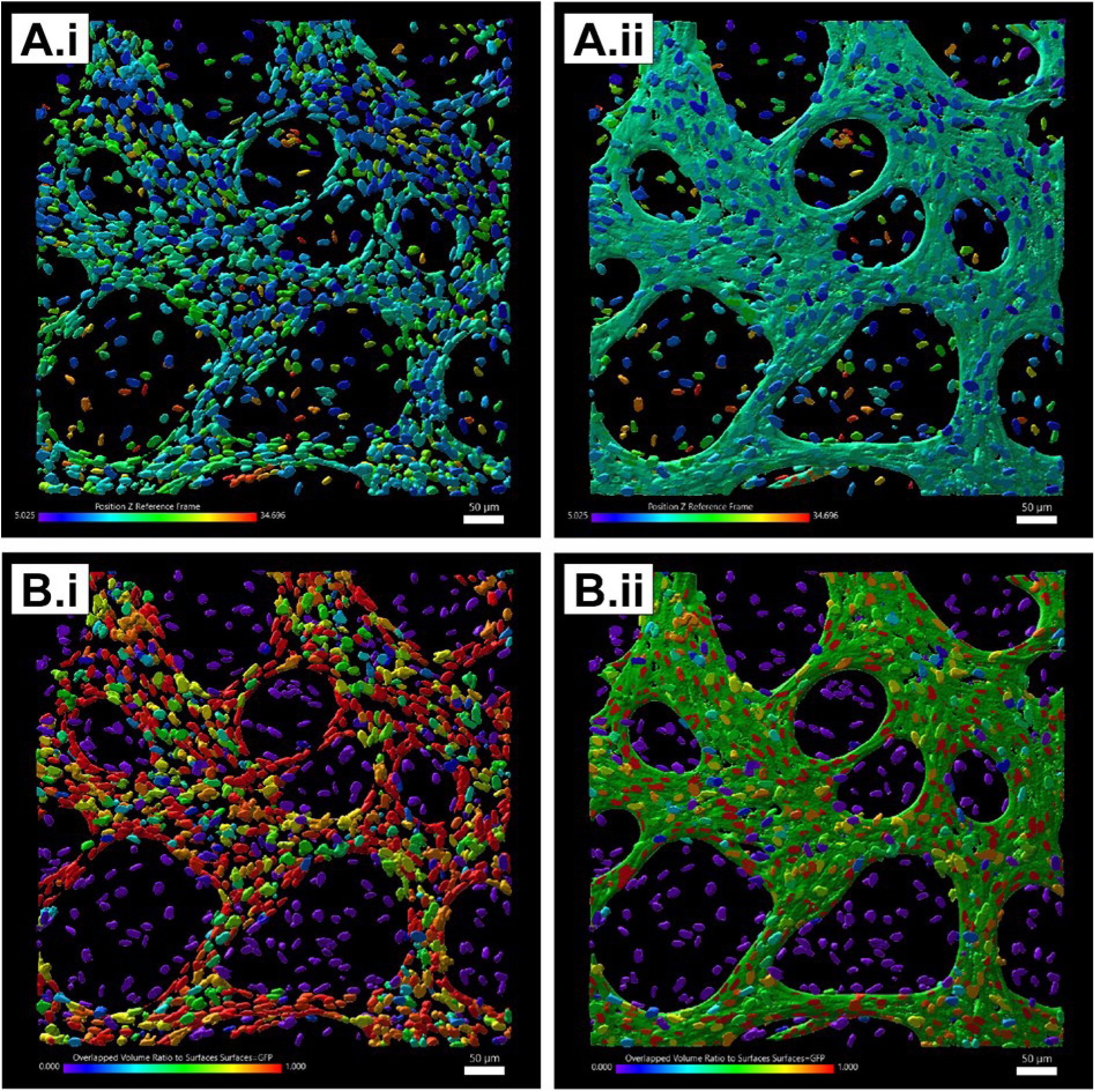
Cell nuclei occupy extra-vascular space between, above and below microvascular networks. **A.** Z-position depth map depicting the height (µm) of segmented cell nuclei within the 582 × 582 × 35.6 µm field of view displayed in the absence (i) and presence of segmented microvascular network GFP (scale bars = 50 µm). **B.** Overlap map depicting the percentage overlap of a cell nuclei with the segmented microvascular network GFP, displayed in the absence (i) and presence of segmented microvascular network GFP (scale bars = 50 µm).

**Figure S14.**
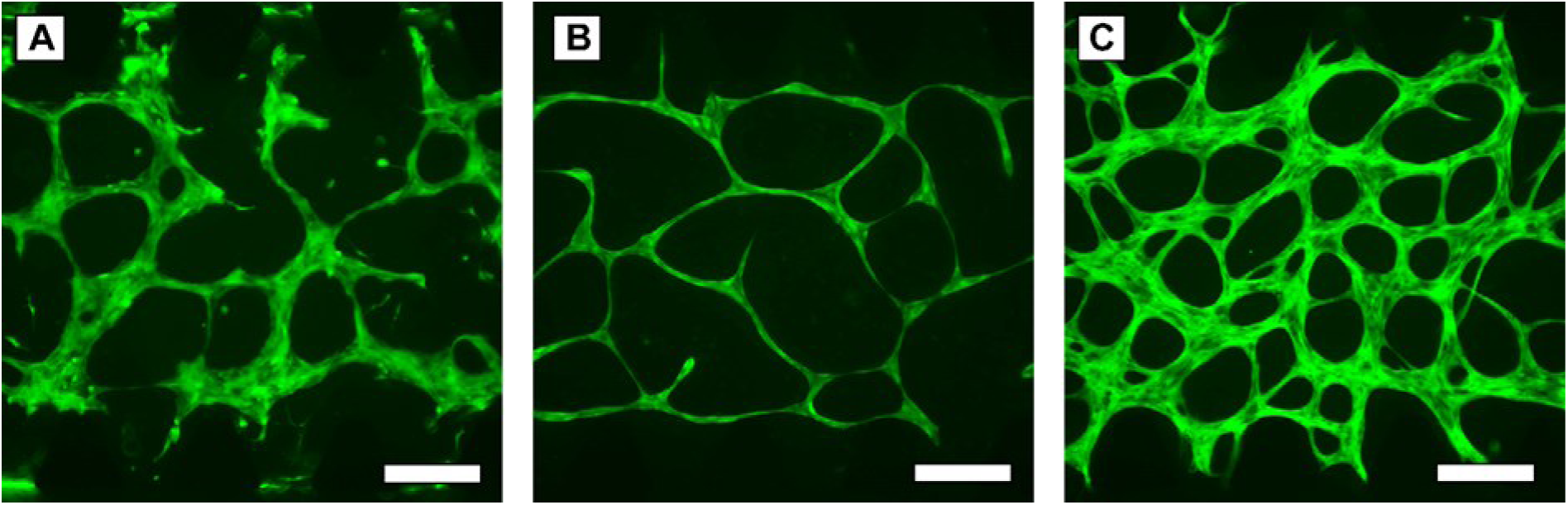
Vessel networks form in altered culture conditions with variable quality. **A.** In the absence of adipose-derived mesenchymal stem cells (hAD-MSCs) vessel networks form with discontinuities and rough borders (scale bar = 300 µm). **B.** In the absence of vascular cell growth factors (in the presence of hAD-MSCs, 1:1) continuous vessel networks with tight borders and abnormally thin vessel diameters form (scale bar = 300 µm). **C.** In the presence of both hAD-MSCs (1:1 seeding ratio) and vascular cell growth factors continuous vessel networks with tight borders and typical vessel diameters form (scale bar = 300 µm).

**Figure S15.**
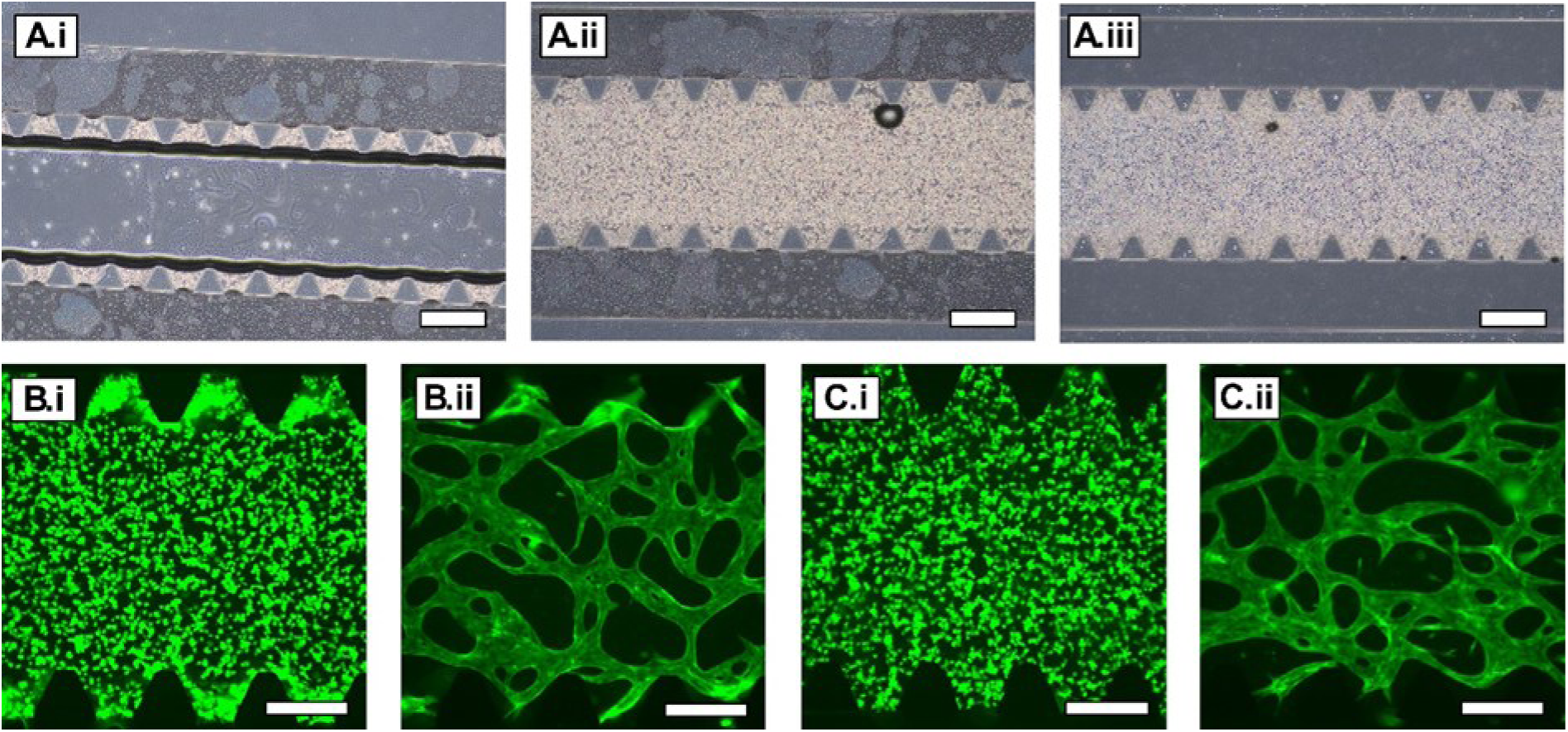
An alternative seeding strategy improved inter-pillar vessels but does not support network opening. **A.** A 100% suspension of 13 × 10^6^ GFP-hAECs/mL in hydrogel precursor is loaded into the culture compartment and immediately aspirated leaving cell suspension between pillars (i). The remainder of the culture compartment is then filled with a 1:1 GFP-hAEC:hAD-MSC hydrogel precursor suspension (ii) and gelled for 15 minutes and then channels filled with complete vascular cell media (iii, scale bars = 500 µm). **B.** Green fluorescence images of the alternate seeding strategy at days 0 (i) and 7 (ii, scale bar = 300 µm). **C.** Green fluorescence images of the typical seeding strategy at days 0 (i) and 7 (ii, scale bar = 300 µm).

**Figure S16.**
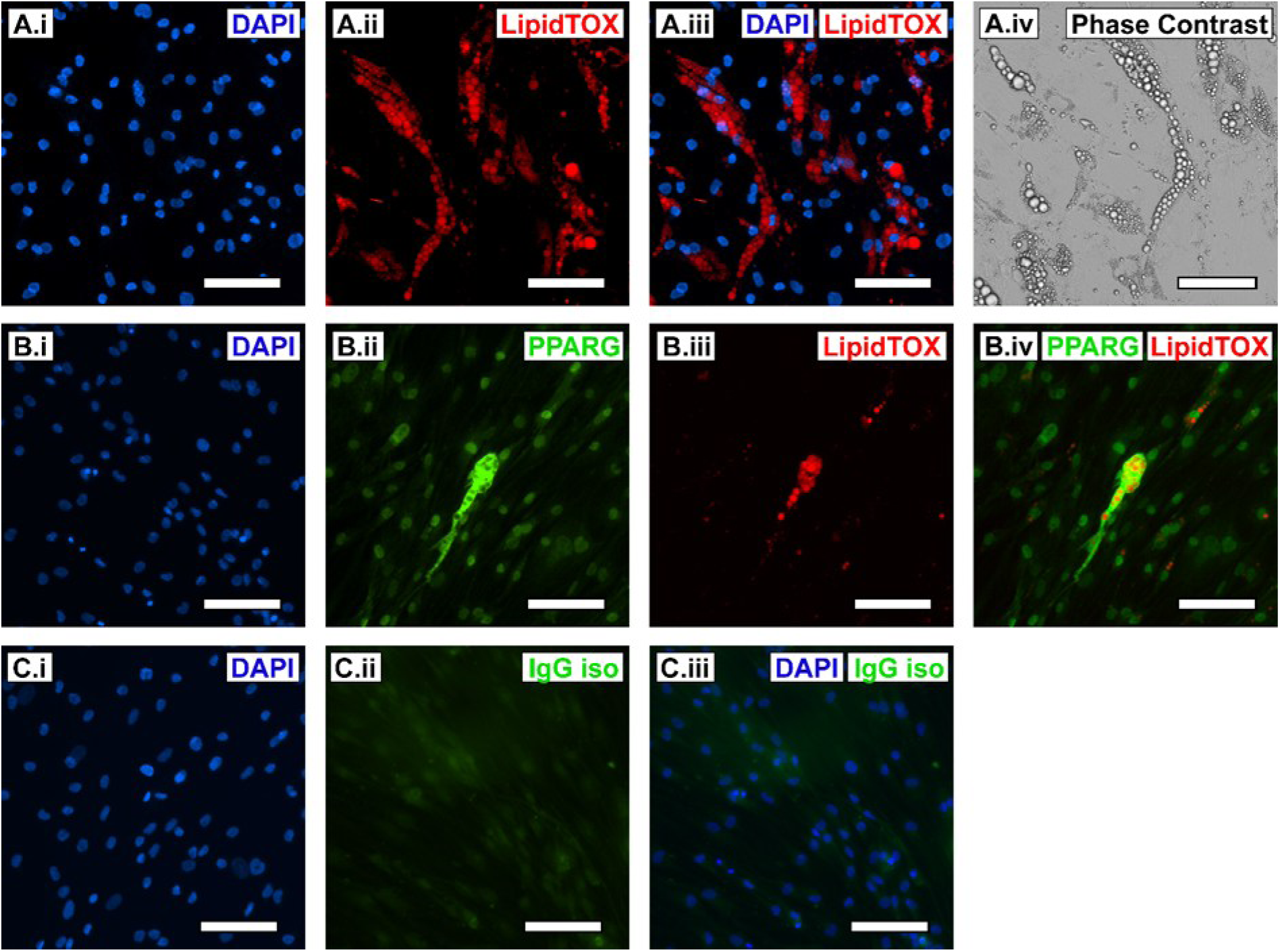
hAD-MSCs exhibit a characteristic adipocyte phenotype after 17 days of 2D differentiation culture. **A.** Lipid-sensitive fluorescent LipidTOX stains lipid droplets, panels depict DAPI nuclear counterstain (i), LipidTOX (ii), DAPI-LipidTOX merge (iii) and phase contrast. **B.** Immunocytochemical detection for peroxisome proliferator-activated receptor gamma (PPARG), panels depict DAPI nuclear counterstain (i), PPARG (ii), LipidTOX (iii) and PPARG- LipidTOX merge (iv). **C.** Rabbit IgG isotype control, panels depict DAPI nuclear counter stain (i), IgG isotype control antibody (ii) and DAPI-IgG isotype control antibody merge (iii). Scale bars = 100 µm.

## Supplemental Material for Growth Factor Distribution Modelling

Simplified code used to generate growth factor distribution models and plots like those found **figure 3**. Code has been simplified to include a single culture media change where vascular growth factor IGF-1 is added from left-hand top and bottom media channels and stromal growth factor EGF is added from right-hand top and bottom channels. The simulations create a 3D matrix of each growth factor, representing its distribution across the hydrogel culture compartment (*xy*) and time. The 3D matrices are then plotted in three ways similar as represented in **figure 3.C**, a 3D surface plot of maximal growth factor concentrations for each *xy* location in culture, a 2D line plot of average relative growth factor concentration over the length of the culture compartment over time, and a 2D heatmap of maximal relative growth factor concentrations. The second two plots have superimposed lines relating to the position where the half-maximal effective concentration of each growth factor is reached across the culture compartment.

Code can be run in Python, dependent on numpy and matplotlib.pyplot packages.

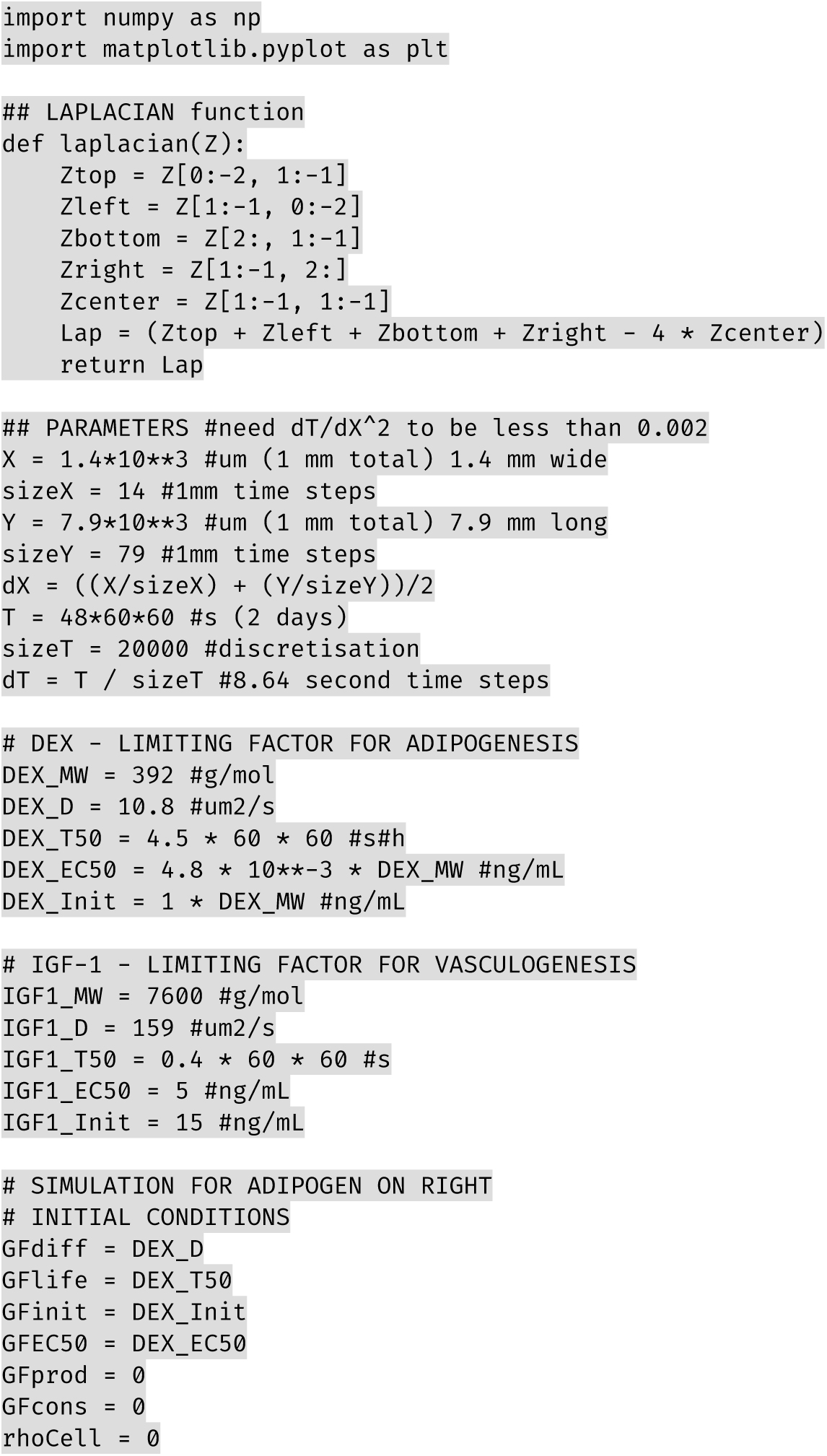
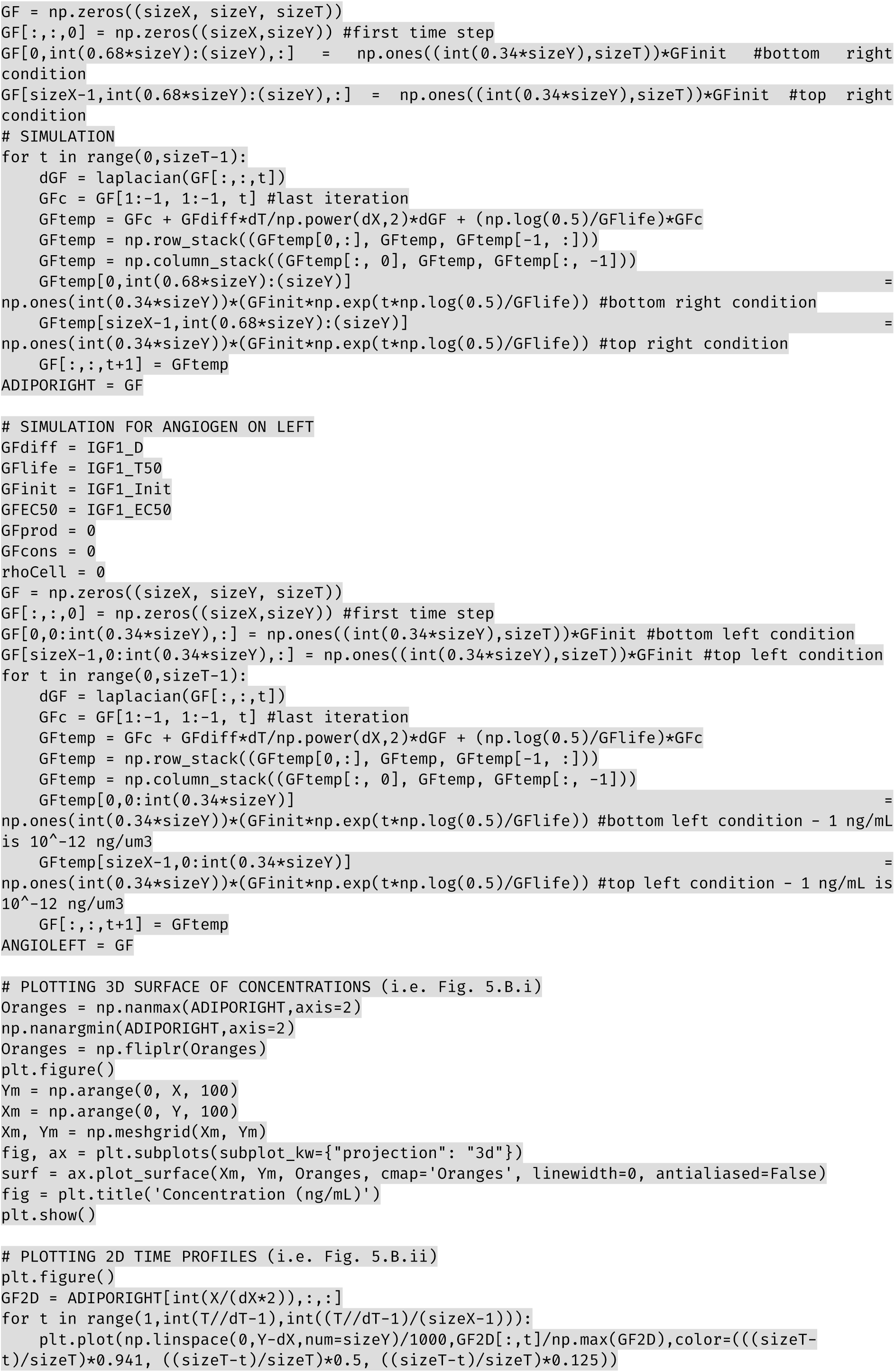
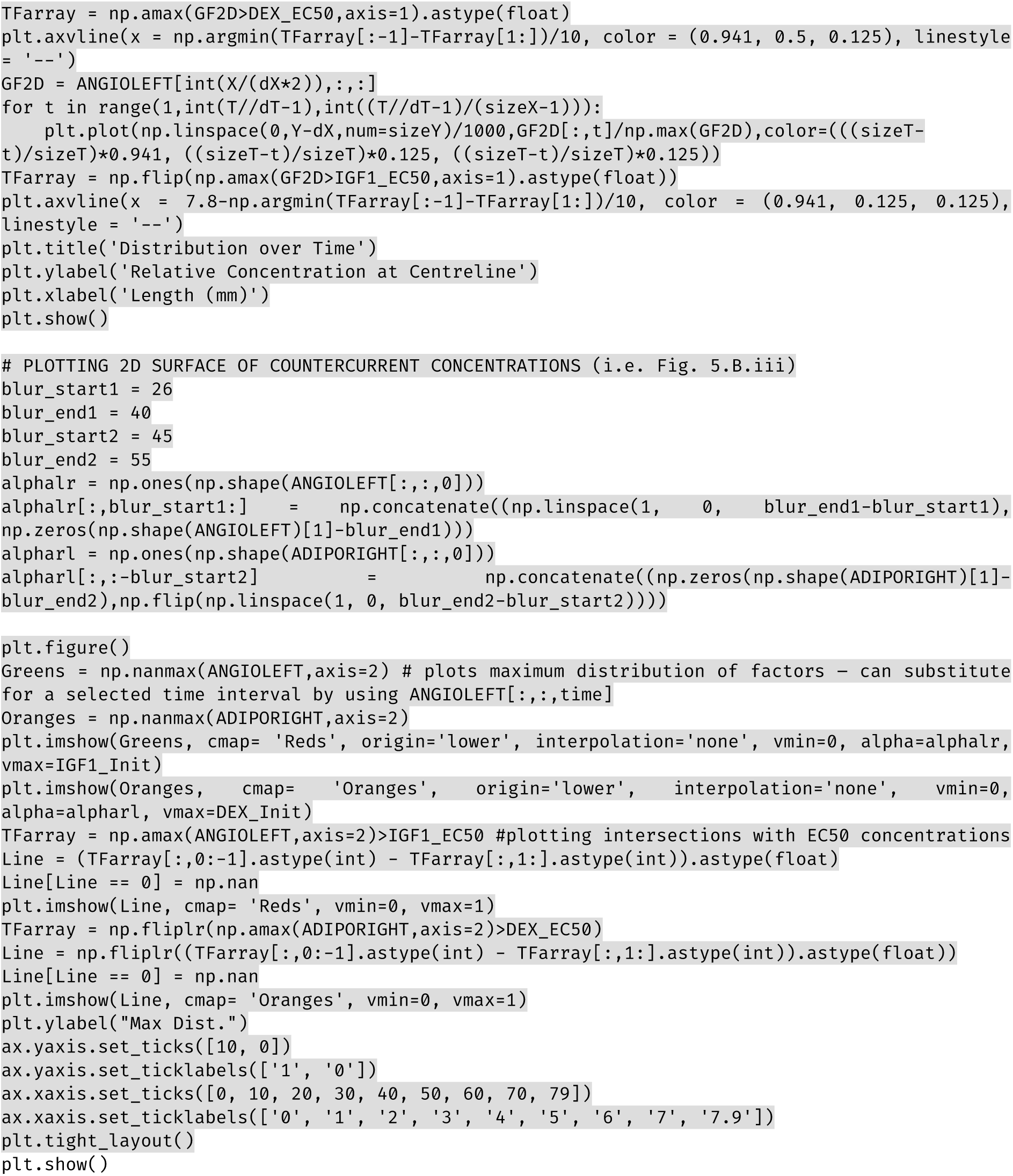

**Figure S17.**
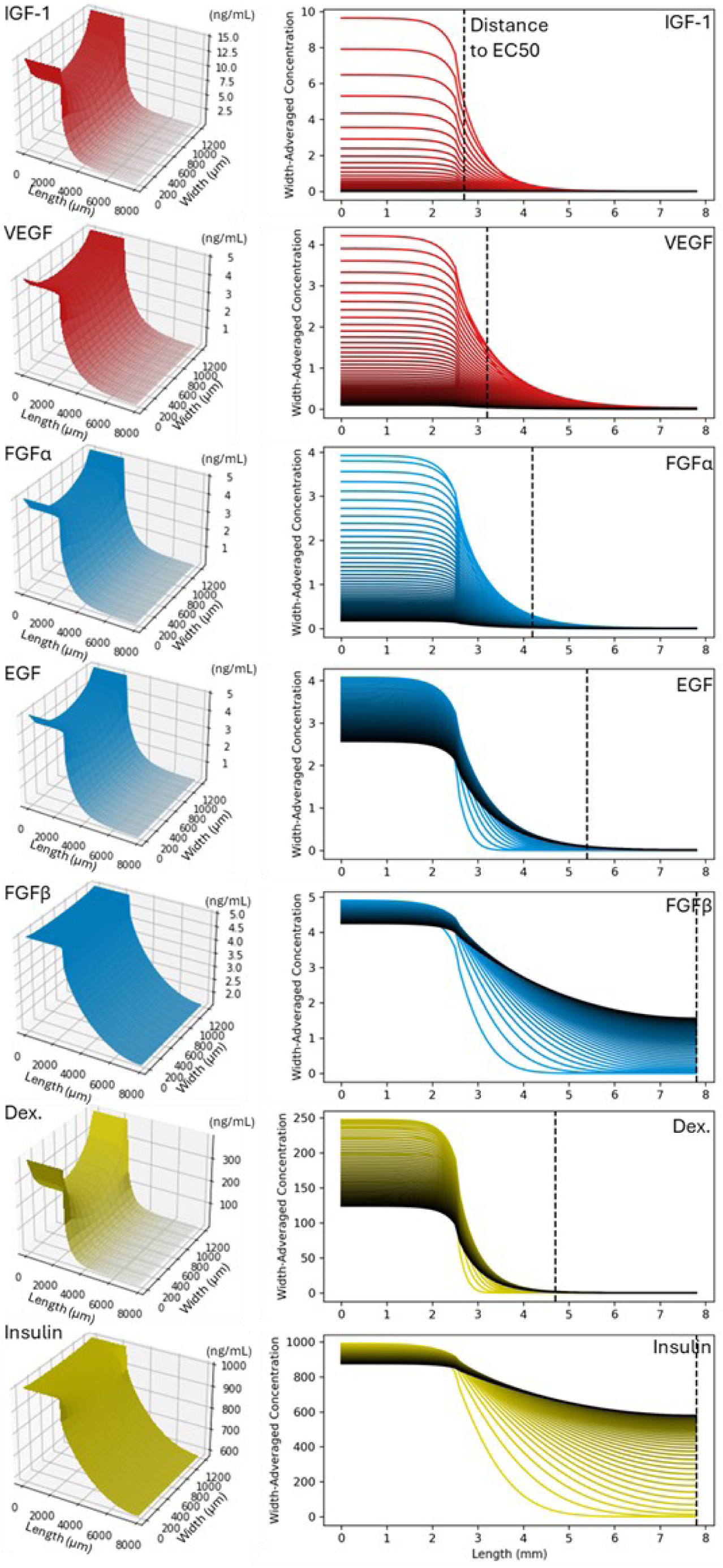
Simulated gradients of stromal maintenance, vasculogenic and adipogenic growth factors. The vascular (Vasc, red), stromal (MSC, blue), adipogenic (Adipo, yellow) growth factors IGF-1, VEGF, FGF-α, EGF, FGF-β, DEX andinsulin. The maximum simulated concentration (in ng/mL) of these factors for each mm^2^ of culture compartment area (13 × 79 mm) over 2 days (i). Growth factor concentrations across the culture compartment length at mid-width, relative to supplemented concentrations, where line colour turns darker for each hour of culture up to 2 days (ii). IGF-1: insulin-like growth factor 1, VEGF: vascular endothelial growth factor, FGF-α: fibroblast growth factor acidic, EGF: epidermal growth factor, FGF-β: fibroblast growth factor basic, DEX: dexamethasone.

**Figure S18.**
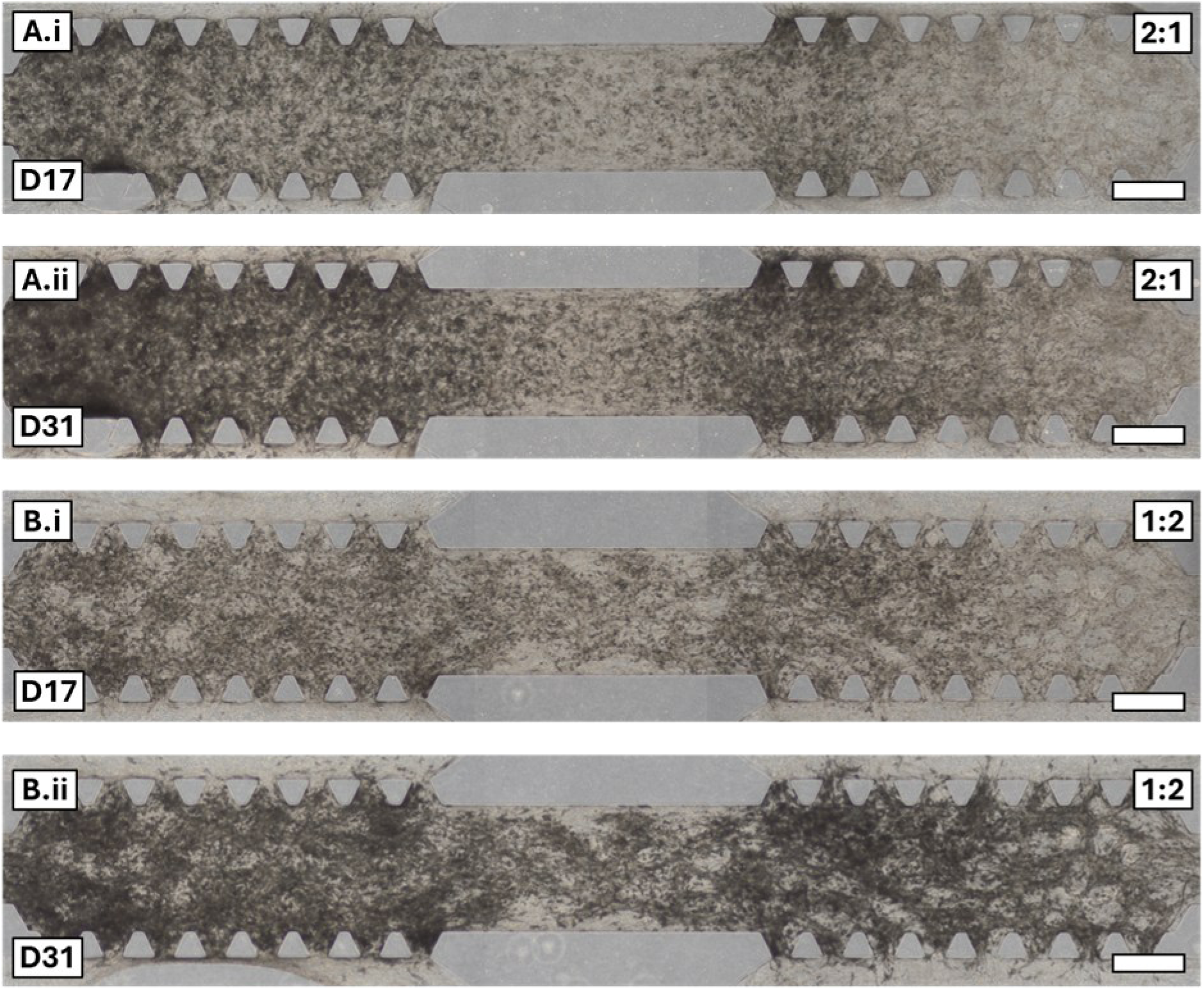
GFP-hAEC) and hAD-MSC gradient co-culture supports the co-formation of microvascular networks and adipogenic differentiation. A. Colour brightfield microscopy images of 2:1 seeding ratio microfluidic device cultures after 17 (i) and 31-days (ii, scale bars = 500 µm). B. Colour brightfield microscopy images of 1:2 seeding ratio microfluidic device cultures after 17 (i) and 31-days (ii, scale bars = 500 µm).

**Figure S19.**
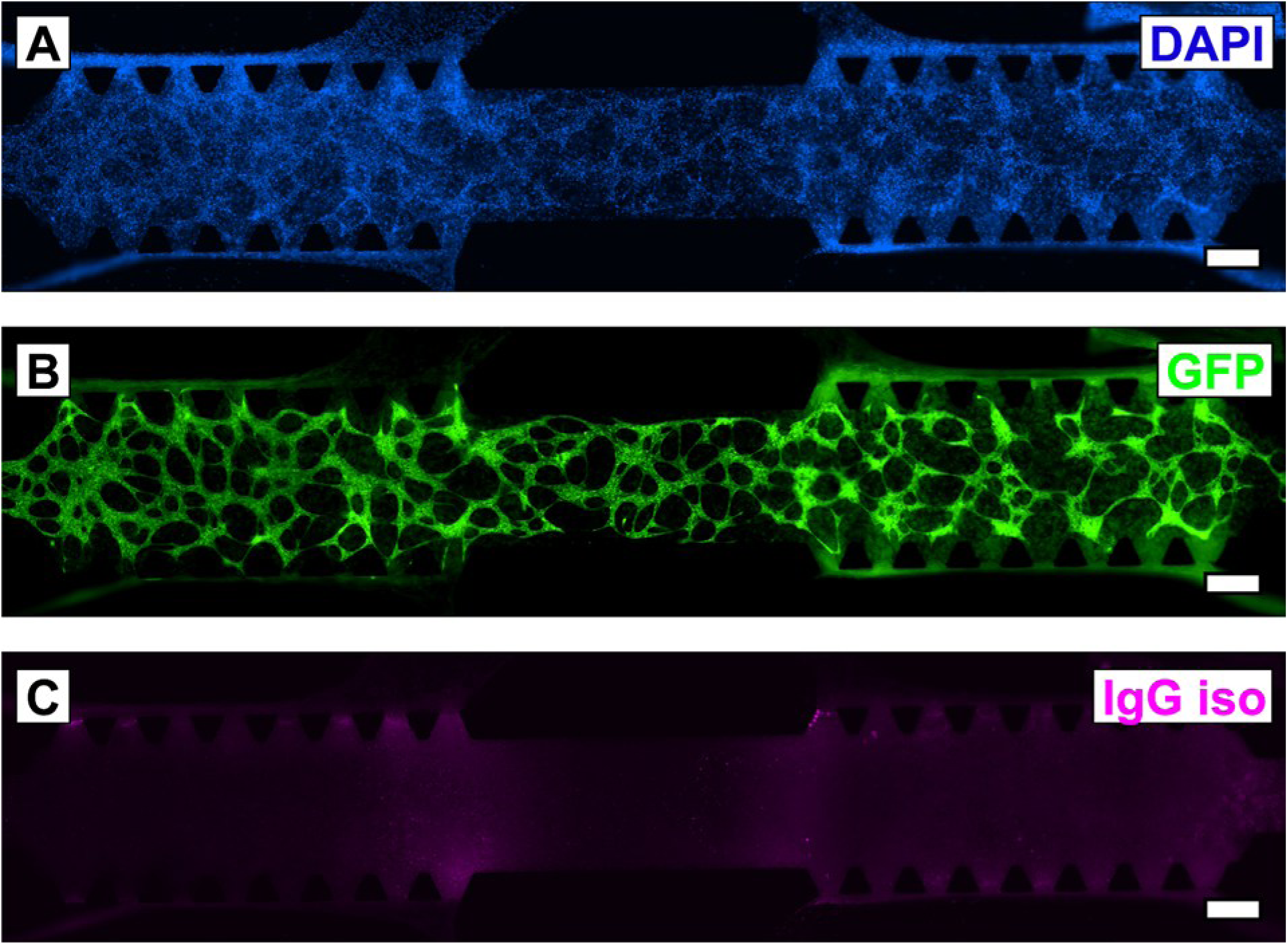
Immunocytochemical detection of 1:1 co-cultures at day 17 for IgG isotype control antibody. Fluorescence mages depicting the nuclear counter stain DAPI (A), GFP (B) and rabbit IgG isotype control (C, scale bars = 500 µm).

**Figure S20.**
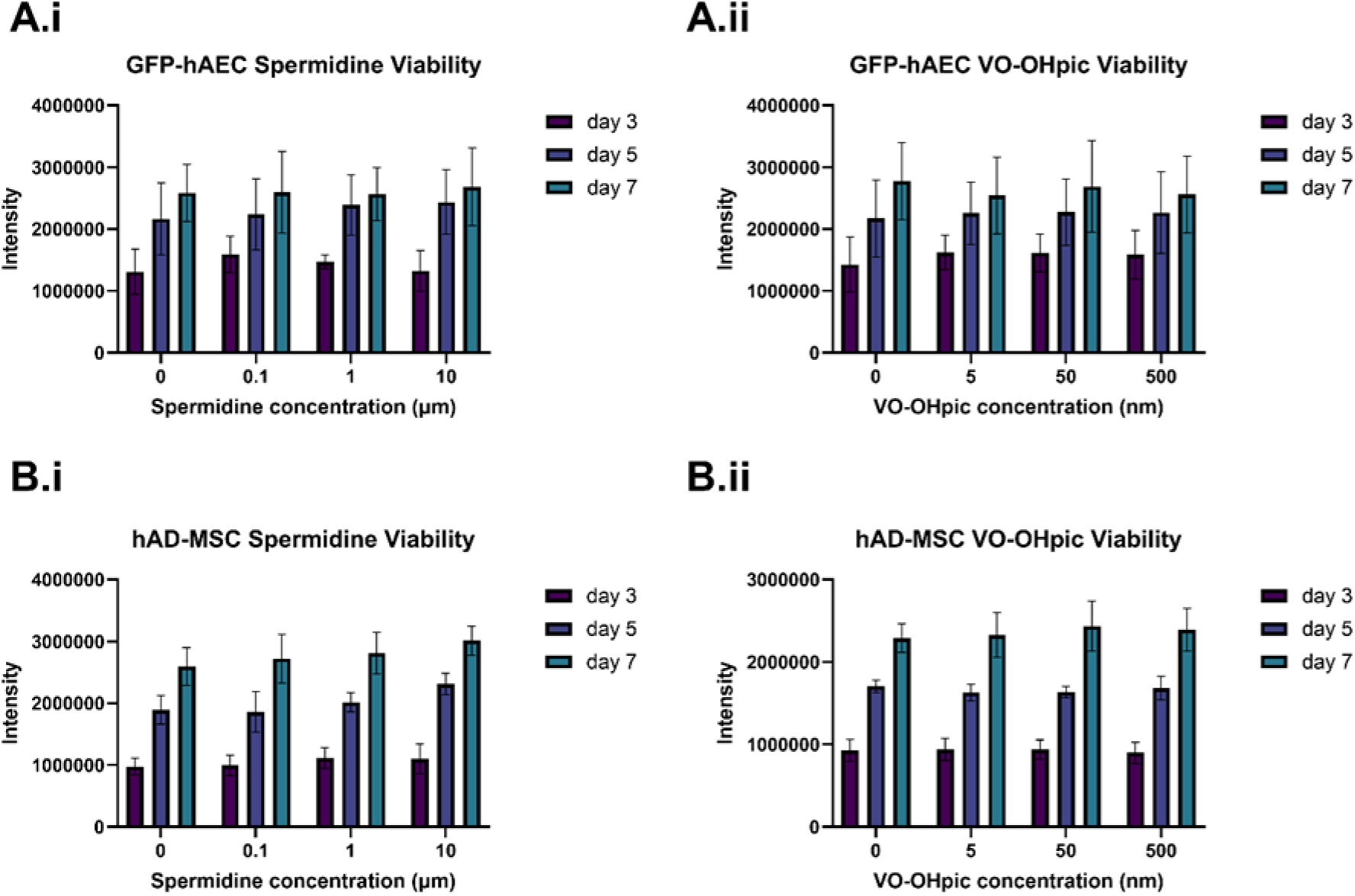
Both spermidine and hydroxyl(oxo)vanadium 3-hydroxypiridine-2-carboxylic acid (VO-OHpic) exhibit no significant influence on GFP-hAEC or hAD-MSC mono-cultures. **A.** GFP-hAECs cultured for 3, 5 and 7-days in media supplemented with 0–10 µM spermidine (i) and 0–500 nM VO-OHpic (ii) (N = 3 biological replicas, mean ± standard deviation). Results were compared via an ordinary two-way ANOVA followed by a Tukey multiple comparisons test. **B.** hAD-MSCs cultured for 3, 5 and 7-days in media supplemented with 0–10 µM spermidine (i) and 0–500 nM VO-OHpic (ii) (N = 3 biological replicas, mean ± standard deviation). Results were compared via an ordinary two-way ANOVA followed by a Tukey multiple comparisons test.

**Figure S21.**
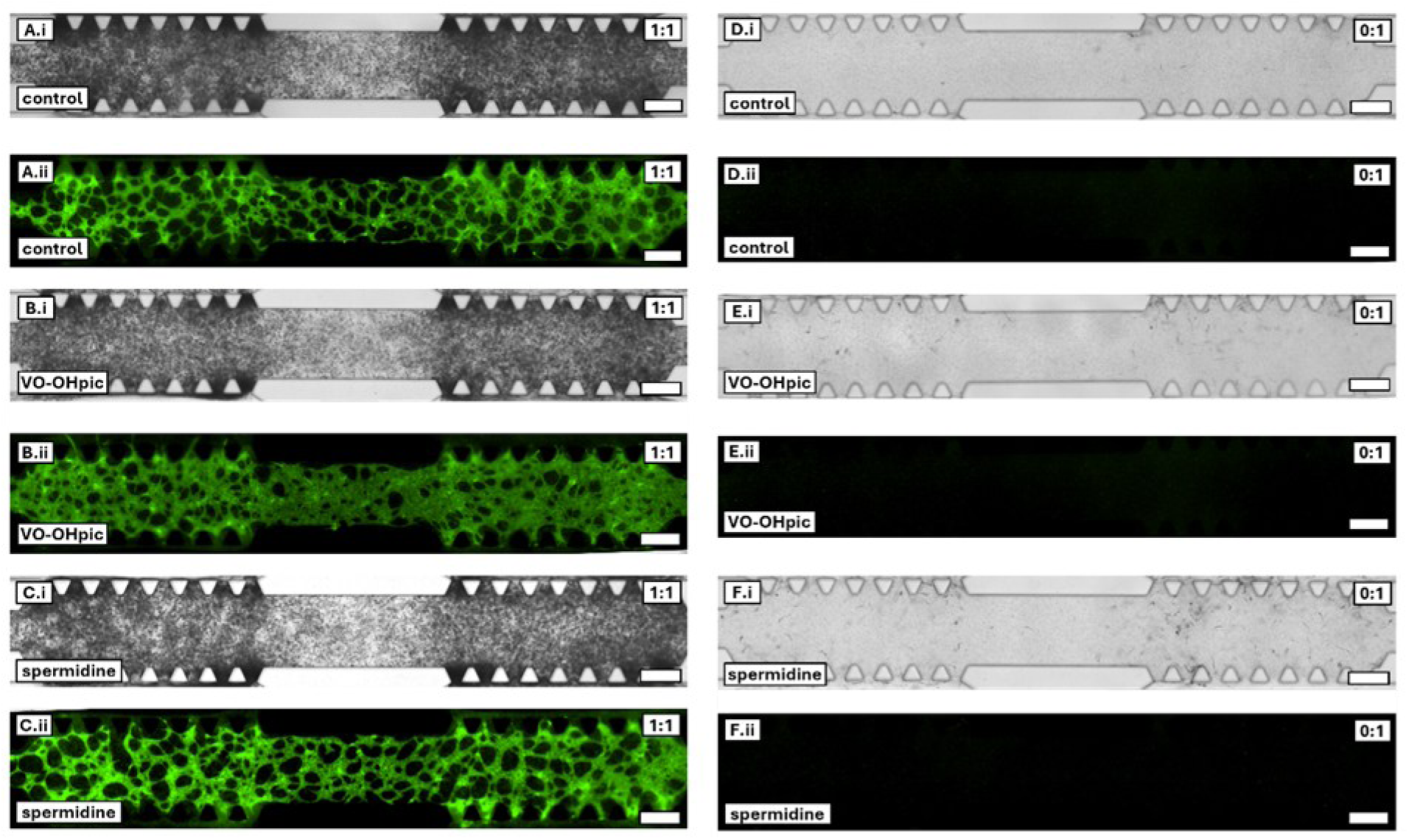
Inducers of lipolysis do not inhibit the formation of lipids in GFP-hAEC and hAD- MSC gradient co-cultures. Monochromatic brightfield (i) and green fluorescence (ii) images of day 31 1:1 seeding ratio cultures without lipolytic supplements (**A**) with 50 nM VO-OHpic (**B**) and 1 µM spermidine (**C**, scale bars = 500 µm). Monochromatic brightfield (i) and green fluorescence (ii) images of day 31 0:1 seeding ratio cultures without lipolytic supplements (**D**) with 50 nM VO- OHpic (**E**) and 1 µM spermidine (**F**, scale bars = 500 µm).

## References

1. Ajmal, L., S. Ajmal, M. Ajmal, and G. Nawaz, Organ Regeneration Through Stem Cells and Tissue Engineering. Cureus, 2023. 15(1): p. e34336.

2. Rouwkema, J., et al., Supply of nutrients to cells in engineered tissues. Biotechnol Genet Eng Rev, 2010. 26: p. 163–78.

3. Luk, C., N.J. Haywood, K.I. Bridge, and M.T. Kearney, Paracrine Role of the Endothelium in Metabolic Homeostasis in Health and Nutrient Excess. Front Cardiovasc Med, 2022. 9: p. 882923.

4. Monelli, E., et al., Angiocrine polyamine production regulates adiposity. Nature Metabolism, 2022. 4(3): p. 327–343.

5. Christov, C., et al., Muscle Satellite Cells and Endothelial Cells: Close Neighbors and Privileged Partners. Molecular Biology of the Cell, 2007. 18(4): p. 1397–1409.

6. He, S., et al., Endothelial POFUT1 controls injury-induced liver fibrosis by repressing fibrinogen synthesis. Journal of Hepatology, 2024. 81(1): p. 135–148.

7. Rafii, S., J.M. Butler, and B.-S. Ding, Angiocrine functions of organ-specific endothelial cells. Nature, 2016. 529(7586): p. 316-325.

8. Rademakers, T., J.M. Horvath, C.A. van Blitterswijk, and V.L.S. LaPointe, Oxygen and nutrient delivery in tissue engineering: Approaches to graft vascularization. J Tissue Eng Regen Med, 2019. 13(10): p. 1815–1829.

9. Jensen, C. and Y. Teng, Is It Time to Start Transitioning From 2D to 3D Cell Culture? Front Mol Biosci, 2020. 7: p. 33.

10. Vis, M.A.M., K. Ito, and S. Hofmann, Impact of Culture Medium on Cellular Interactions in in vitro Co-culture Systems. Frontiers in Bioengineering and Biotechnology, 2020. 8.

11. Langhans, S.A., Three-Dimensional in Vitro Cell Culture Models in Drug Discovery and Drug Repositioning. Frontiers in Pharmacology, 2018. 9.

12. Chung, B.G. and J. Choo, Microfluidic gradient platforms for controlling cellular behavior. Electrophoresis, 2010. 31(18): p. 3014–27.

13. Murphy, A.R. and M.C. Allenby, In vitro microvascular engineering approaches and strategies for interstitial tissue integration. Acta Biomaterialia, 2023. 171: p. 114–130.

14. Ceddia, R.B., The role of AMP-activated protein kinase in regulating white adipose tissue metabolism. Molecular and Cellular Endocrinology, 2013. 366(2): p. 194–203.

15. Corvera, S., Cellular Heterogeneity in Adipose Tissues. Annual Review of Physiology, 2021. 83(Volume 83, 2021): p. 257-278.

16. Dufau, J., et al., In vitro and ex vivo models of adipocytes. Am J Physiol Cell Physiol, 2021. 320(5): p. C822–c841.

17. Ioannidou, A., R.M. Fisher, and C.E. Hagberg, The multifaceted roles of the adipose tissue vasculature. Obes Rev, 2022. 23(4): p. e13403.

18. Louis, F., et al., Injectable Prevascularized Mature Adipose Tissues (iPAT) to Achieve Long-Term Survival in Soft Tissue Regeneration. Advanced Healthcare Materials, 2022. 11(23): p. 2201440.

19. Struss, M. and E. Bellas, Microphysiological Modeling of Vascular Adipose Tissue for Multi-Throughput Applications. bioRxiv, 2024: p. 2024.01.30.578061.

20. Zhou, Y., H. Li, and N. Xia, The Interplay Between Adipose Tissue and Vasculature: Role of Oxidative Stress in Obesity. Front Cardiovasc Med, 2021. 8: p. 650214.

21. Corvera, S. and O. Gealekman, Adipose tissue angiogenesis: impact on obesity and type-2 diabetes. Biochim Biophys Acta, 2014. 1842(3): p. 463–72.

22. Polkinghorne, M.D., H.W. West, and C. Antoniades, Adipose Tissue in Cardiovascular Disease: From Basic Science to Clinical Translation. Annual Review of Physiology, 2024. 86(Volume 86, 2024): p. 175-198.

23. James, M., T.P. Varghese, R. Sharma, and S. Chand, Association Between Metabolic Syndrome and Diabetes Mellitus According to International Diabetic Federation and National Cholesterol Education Program Adult Treatment Panel III Criteria: a Cross-sectional Study. J Diabetes Metab Disord, 2020. 19(1): p. 437–443.

24. Hammel, J.H. and E. Bellas, Endothelial cell crosstalk improves browning but hinders white adipocyte maturation in 3D engineered adipose tissue. Integrative Biology, 2020. 12(4): p. 81–89.

25. Xu, M., et al., An cell-assembly derived physiological 3D model of the metabolic syndrome, based on adipose-derived stromal cells and a gelatin/alginate/fibrinogen matrix. Biomaterials, 2010. 31(14): p. 3868–3877.

26. Evensen, L., et al., Mural Cell Associated VEGF Is Required for Organotypic Vessel Formation. PLOS ONE, 2009. 4(6): p. e5798.

27. Andrée, B., et al., Formation of three-dimensional tubular endothelial cell networks under defined serum-free cell culture conditions in human collagen hydrogels. Scientific Reports, 2019. 9(1): p. 5437.

28. Wan, Z., et al., A Robust Method for Perfusable Microvascular Network Formation In Vitro. Small Methods, 2022. 6(6): p. e2200143.

29. Zudaire, E., L. Gambardella, C. Kurcz, and S. Vermeren, A Computational Tool for Quantitative Analysis of Vascular Networks. PLOS ONE, 2011. 6(11): p. e27385.

30. Schindelin, J., et al., Fiji: an open-source platform for biological-image analysis. Nature Methods, 2012. 9(7): p. 676–682.

31. Altan-Bonnet, G. and R. Mukherjee, Cytokine-mediated communication: a quantitative appraisal of immune complexity. Nature Reviews Immunology, 2019. 19(4): p. 205–217.

32. Allenby, M.C., et al., A Spatiotemporal Microenvironment Model to Improve Design of a Three-Dimensional Bioreactor for Red Cell Production. Tissue Eng Part A, 2022. 28(1-2): p. 38–53.

33. American Type Culture Collection. A Chemically-induced Method of Adipogenesis. 2024; Available from: https://www.atcc.org/resources/technical-documents/chemically-induced-method-of-adipogenesis.

34. Leonidakis, K.A., et al., Fibrin structural and diffusional analysis suggests that fibers are permeable to solute transport. Acta Biomater, 2017. 47: p. 25–39.

35. Le, K.N., et al., Vascular regeneration by local growth factor release is self-limited by microvascular clearance. Circulation, 2009. 119(22): p. 2928–35.

36. BailÓN-Plaza, A. and M.C.H. Van Der Meulen, A Mathematical Framework to Study the Effects of Growth Factor Influences on Fracture Healing. Journal of Theoretical Biology, 2001. 212(2): p. 191–209.

37. Xia, X., et al., Pharmacokinetic Properties of 2nd-Generation Fibroblast Growth Factor-1 Mutants for Therapeutic Application. PLOS ONE, 2012. 7(11): p. e48210.

38. Culajay, J.F., S.I. Blaber, A. Khurana, and M. Blaber, Thermodynamic Characterization of Mutants of Human Fibroblast Growth Factor 1 with an Increased Physiological Half-Life. Biochemistry, 2000. 39(24): p. 7153–7158.

39. STEMCELL Technologies. Human Recombinant FGF-acidic. 2024; Available from: https://www.stemcell.com/products/human-recombinant-fgf-acidic.html.

40. Zhang, P. and C. Liu, Enhancement of Skin Wound Healing by rhEGF-Loaded Carboxymethyl Chitosan Nanoparticles. Polymers, 2020. 12(7): p. 1612.

41. Johnston, S.T., et al., Estimating cell diffusivity and cell proliferation rate by interpreting IncuCyte ZOOM™ assay data using the Fisher-Kolmogorov model. BMC Systems Biology, 2015. 9(1): p. 38.

42. Kuo, C.-Y., et al., Development of a 3D Printed, Bioengineered Placenta Model to Evaluate the Role of Trophoblast Migration in Preeclampsia. ACS Biomaterials Science & Engineering, 2016. 2(10): p. 1817–1826.

43. STEMCELL Technologies. Human Recombinant EGF. 2024; Available from: https://www.stemcell.com/products/human-recombinant-egf.html.

44. Thorne, R.G., S. Hrabetová, and C. Nicholson, Diffusion of epidermal growth factor in rat brain extracellular space measured by integrative optical imaging. J Neurophysiol, 2004. 92(6): p. 3471–81.

45. Andasari, V., et al., Computational model of wound healing: EGF secreted by fibroblasts promotes delayed re-epithelialization of epithelial keratinocytes. Integrative Biology, 2018. 10(10): p. 605–634.

46. Ding, I. and A.M. Peterson, Half-life modeling of basic fibroblast growth factor released from growth factor-eluting polyelectrolyte multilayers. Scientific Reports, 2021. 11(1): p. 9808.

47. Filion, R.J. and A.S. Popel, A Reaction-Diffusion Model of Basic Fibroblast Growth Factor Interactions with Cell Surface Receptors. Annals of Biomedical Engineering, 2004. 32(5): p. 645–663.

48. STEMCELL Technologies. Human Recombinant bFGF. 2024; Available from: https://www.stemcell.com/products/human-recombinant-bfgf.html.

49. Nauman, J.V., P.G. Campbell, F. Lanni, and J.L. Anderson, Diffusion of insulin-like growth factor-I and ribonuclease through fibrin gels. Biophys J, 2007. 92(12): p. 4444–50.

50. Hellstrom, A., et al., IGF-1 as a Drug for Preterm Infants: A Step-Wise Clinical Development. Current Pharmaceutical Design, 2017. 23(38): p. 5964–5970.

51. Simón-Yarza, T., et al., Vascular endothelial growth factor-delivery systems for cardiac repair: an overview. Theranostics, 2012. 2(6): p. 541–52.

52. Miura, T. and R. Tanaka, In vitro Vasculogenesis Models Revisited - Measurement of VEGF Diffusion in Matrigel. Math. Model. Nat. Phenom., 2009. 4(4): p. 118–130.

53. Bedell, W. and A.D. Stroock, A VEGF reaction-diffusion mechanism that selects variable densities of endothelial tip cells. bioRxiv, 2019: p. 624999.

54. Baker, Q.B., G.J. Podgorski, E. Vargis, and N.S. Flann, A computational study of VEGF production by patterned retinal epithelial cell colonies as a model for neovascular macular degeneration. Journal of Biological Engineering, 2017. 11(1): p. 26.

55. Kim, J. and A. Chauhan, Dexamethasone transport and ocular delivery from poly(hydroxyethyl methacrylate) gels. International Journal of Pharmaceutics, 2008. 353(1): p. 205–222.

56. Sidibeh, C.O., et al., FKBP5 expression in human adipose tissue: potential role in glucose and lipid metabolism, adipogenesis and type 2 diabetes. Endocrine, 2018. 62(1): p. 116–128.

57. Mooradian, A.D. and C.N. Mariash, Effects of Insulin and Glucose on Cultured Rat Hepatocyte Gene Expression. Diabetes, 1987. 36(8): p. 938–943.

58. Shorten, P.R., C.D. McMahon, and T.K. Soboleva, Insulin Transport within Skeletal Muscle Transverse Tubule Networks. Biophysical Journal, 2007. 93(9): p. 3001–3007.

59. Weiland, M., et al., The signaling potential of the receptors for insulin and insulin-like growth factor I (IGF-I) in 3T3-L1 adipocytes: comparison of glucose transport activity, induction of oncogene c-fos, glucose transporter mRNA, and DNA-synthesis. J Cell Physiol, 1991. 149(3): p. 428–35.

60. Rossant, C., *Simulating* a partial differential equation — reaction-diffusion systems and Turing patterns, in IPython Interactive Computing and Visualization Cookbook: Over 100 hands-on recipes to sharpen your skills in high-performance numerical computing and data science in the Jupyter Notebook. 2018, Packt Publishing Ltd.

61. Kim, S., H. Lee, M. Chung, and N.L. Jeon, Engineering of functional, perfusable 3D microvascular networks on a chip. Lab on a Chip, 2013. 13(8): p. 1489–1500.

62. Campisi, M., et al., 3D self-organized microvascular model of the human blood-brain barrier with endothelial cells, pericytes and astrocytes. Biomaterials, 2018. 180: p. 117–129.

63. Nowwarote, N., et al., PTEN regulates proliferation and osteogenesis of dental pulp cells and adipogenesis of human adipose-derived stem cells. Oral Dis, 2023. 29(2): p. 735–746.

64. Bi, M., et al., Cell-based mechanisms and strategies of co-culture system both in vivo and vitro for bone tissue engineering. Biomedicine & Pharmacotherapy, 2023. 169: p. 115907.

65. Chan, G., et al., In vivo optical imaging of human retinal capillary networks using speckle variance optical coherence tomography with quantitative clinico-histological correlation. Microvascular Research, 2015. 100: p. 32–39.

66. Pusztaszeri, M.P., W. Seelentag, and F.T. Bosman, Immunohistochemical expression of endothelial markers CD31, CD34, von Willebrand factor, and Fli-1 in normal human tissues. J Histochem Cytochem, 2006. 54(4): p. 385–95.

67. Armulik, A., G. Genové, and C. Betsholtz, Pericytes: Developmental, Physiological, and Pathological Perspectives, Problems, and Promises. Developmental Cell, 2011. 21(2): p. 193–215.

68. Wang, Y., et al., Relative contributions of adipose-resident CD146(+) pericytes and CD34(+) adventitial progenitor cells in bone tissue engineering. NPJ Regen Med, 2019. 4:p. 1.

69. Moraes, D.A., et al., A reduction in CD90 (THY-1) expression results in increased differentiation of mesenchymal stromal cells. Stem Cell Research & Therapy, 2016. 7(1): p. 97.

70. Venkatakrishnan, G. and V.D. Parvathi, Decoding the mechanism of vascular morphogenesis to explore future prospects in targeted tumor therapy. Medical Oncology, 2022. 39(11): p. 178.

71. Feng, J. and Z. Meng, Insulin growth factor-1 promotes the proliferation and osteogenic differentiation of bone marrow mesenchymal stem cells through the Wnt/β-catenin pathway. Exp Ther Med, 2021. 22(2): p. 891.

72. Hajal, C., et al., Engineered human blood–brain barrier microfluidic model for vascular permeability analyses. Nature Protocols, 2022. 17(1): p. 95–128.

73. Coffin, S.T. and G.R. Gaudette, Aprotinin extends mechanical integrity time of cell-seeded fibrin sutures. J Biomed Mater Res A, 2016. 104(9): p. 2271–9.

74. Chen, R. and D. Dean, Mechanical properties of stem cells from different sources during vascular smooth muscle cell differentiation. Mol Cell Biomech, 2017. 14(3): p. 153–169.

75. Blobel, C.P., 3D trumps 2D when studying endothelial cells. Blood, 2010. 115(25): p. 5128–5130.

76. Ge, Q., et al., VEGF secreted by mesenchymal stem cells mediates the differentiation of endothelial progenitor cells into endothelial cells via paracrine mechanisms. Mol Med Rep, 2018. 17(1): p. 1667–1675.

77. Wan, Z., et al., A robust vasculogenic microfluidic model using human immortalized endothelial cells and Thy1 positive fibroblasts. Biomaterials, 2021. 276: p. 121032.

78. Khaki, M., et al., Mesenchymal Stem Cells Differentiate to Endothelial Cells Using Recombinant Vascular Endothelial Growth Factor-A. Rep Biochem Mol Biol, 2018. 6(2): p. 144–150.

79. Rupnick, M.A., et al., Adipose tissue mass can be regulated through the vasculature. Proc Natl Acad Sci U S A, 2002. 99(16): p. 10730–5.

80. Masnikov, D., et al., hTERT-immortalized adipose-derived stem cell line ASC52Telo demonstrates limited potential for adipose biology research. Anal Biochem, 2021. 628: p. 114268.

81. Rudnicki, M. and T.L. Haas, Adipose tissue lipolysis controlled by endothelial cells. Nature Reviews Endocrinology, 2022. 18(7): p. 397–398.

82. Gliniak, C.M. and P.E. Scherer, A new signal that shrinks fat. Nature Metabolism, 2022. 4(3): p. 305–307.

83. Pulido, R., PTEN Inhibition in Human Disease Therapy. Molecules, 2018. 23(2): p. 285.

84. Mak, L.H., R. Vilar, and R. Woscholski, Characterisation of the PTEN inhibitor VO-OHpic. J Chem Biol, 2010. 3(4): p. 157–63.

85. Liao, C.Y., et al., The Autophagy Inducer Spermidine Protects Against Metabolic Dysfunction During Overnutrition. J Gerontol A Biol Sci Med Sci, 2021. 76(10): p. 1714–1725.

86. Ni, Y., et al., Spermidine activates adipose tissue thermogenesis through autophagy and fibroblast growth factor 21. The Journal of Nutritional Biochemistry, 2024. 125: p. 109569.

87. Ibrahim, M.M., Subcutaneous and visceral adipose tissue: structural and functional differences. Obes Rev, 2010. 11(1): p. 11–8.

88. Belligoli, A., et al., Characterization of subcutaneous and omental adipose tissue in patients with obesity and with different degrees of glucose impairment. Scientific Reports, 2019. 9(1): p. 11333.

89. Schwarz, C., et al., Effects of Spermidine Supplementation on Cognition and Biomarkers in Older Adults With Subjective Cognitive Decline: A Randomized Clinical Trial. JAMA Netw Open, 2022. 5(5): p. e2213875.

90. Senekowitsch, S., et al., High-Dose Spermidine Supplementation Does Not Increase Spermidine Levels in Blood Plasma and Saliva of Healthy Adults: A Randomized Placebo-Controlled Pharmacokinetic and Metabolomic Study. Nutrients, 2023. 15(8).

